# Long-term isolation and introgression shape the genomic distinctiveness of Rice’s Whale

**DOI:** 10.64898/2026.06.15.732430

**Authors:** Diana Aguilar-Gómez, Jacqueline A. Robinson, Christopher C. Kyriazis, Mallory Kenfield, Sergio Nigenda-Morales, Nicole L. Vollmer, Lynsey Wilcox Talbot, Bernard Y. Kim, Ryan D. Hernandez, Patricia E. Rosel, Phillip A. Morin, Kirk E. Lohmueller

## Abstract

Species’ complex demographic history, including population size changes, isolation and gene flow, shapes patterns of genetic variation and deleterious load. In small and declining populations, understanding these processes is critical for predicting inbreeding depression and extinction risk. The Rice’s whale (*Balaenoptera ricei*) is a newly described baleen whale species resident to the heavily industrialized Gulf of Mexico, with a current abundance estimate of 51 (95 % CI: 20–130) individuals, making it one of the most endangered baleen whales globally. Using whole-genome sequences from 25 individuals, we reconstructed the evolutionary and demographic history of Rice’s whale and assessed its genomic health. Our analyses reinforce its distinctiveness from Bryde’s whales, and suggest that Rice’s whale has persisted as a small and isolated population in the Gulf of Mexico for tens of thousands of years. Despite its long-term small effective population size, genomes show modest impacts of inbreeding, including few long runs of homozygosity. We detected a distinct pulse of introgression ∼350 years ago from a Bryde’s whale–like lineage that resulted in windows of elevated heterozygosity in Rice’s whale, though it did not alter the burden of deleterious variation. Forward simulations indicate that a recent population collapse to ∼100 breeding individuals places the species at high risk of future inbreeding and genomic erosion unless population growth occurs. These findings highlight that while gene flow can increase genetic diversity, demographic recovery is essential to mitigate long-term genomic risks, underscoring the importance of management actions that promote sustained population growth.

## Introduction

We are living through a period of rapid biodiversity loss often described as the sixth mass extinction ^1,2^, driven largely by human activities. For species reduced to small, isolated populations, the combined effects of habitat degradation, industrialization, and demographic decline can rapidly erode genetic diversity and increase extinction risk. In such cases, decisions made within the next few decades may determine whether populations recover or are lost.

Few ecosystems illustrate this more starkly than the Gulf of Mexico, one of the world’s most industrialized marine ecosystems, shaped by decades of oil extraction, commercial shipping, and large-scale environmental disturbance ^3,4^. The Rice’s whale (*Balaenoptera ricei*) is the Gulf’s only resident baleen whale species ^5^, and with an estimated population of 51 individuals (95 % CI: 20–130) ^6,7^, one of the most endangered marine mammals in the world. Its restricted range, critically small population, and exposure to a wide number of anthropogenic threats—including vessel strikes, noise pollution, and oil and gas operations—have heightened the urgency of conservation efforts ^5,8^. In populations of this size, genetic processes are expected to play a disproportionate role in shaping viability ^9,10^, yet despite the urgency, the genomic consequences of the species’ demographic history remain mostly uncharacterized.

The reason for this gap in knowledge is because the Rice’s whale was only recently recognized as a distinct species. First documented by Dale Rice ^11^ in the Gulf of Mexico, thought to be a Bryde’s whale (*Balaenoptera edeni*), Gulf of Mexico individuals were later shown to exhibit 10-13% mitochondrial DNA divergence from Bryde’s whales worldwide ^5,88^. Combined with diagnostic cranial morphology, this evidence prompted formal recognition of *B. ricei* as a new species in 2021 ^5^. The species’ brief taxonomic history means that fundamental questions about its evolutionary origins, population structure, and genomic health have scarcely been asked, let alone answered.

The Rice’s whale’s recognition also emerged from a broader and still-unresolved debate about species boundaries within the Bryde’s whale complex. Historically, a single circumglobal species of tropical baleen whale was recognized ^12^, but molecular and morphological evidence has since revealed substantial diversity within this group. This includes the Eden’s whale (*B. edeni eden*i; generally smaller and restricted to the western Pacific and Indian Oceans) and the Bryde’s whale (*B. edeni brydei*, found across all three major ocean basins); they have long been treated as subspecies, however, both mitochondrial ^5,8,13,14^, and nuclear ^15,16^ genomic data consistently place them in separate clades, and distinct morphological characters have been identified ^17^. Their continued classification as subspecies thus remains a matter of active debate ^14,17–21^.

Here, we analyze 25 whole-genome sequences from Rice’s whales to characterize genetic variation, reconstruct evolutionary and demographic history, assess phylogenetic relationships with closely related baleen whale species, and quantify patterns of genetic load. Together, these genomic analyses reveal the Rice’s whale’s unique and complex evolutionary history, while providing the basis for informed conservation decisions that will determine this species’ fate.

## Results

### Rice’s whale distinctiveness within the baleen whale phylogeny

We combined genomic resequencing data from Rice’s whales with newly generated and publicly available baleen whale genomes and a chromosome-level Rice’s whale assembly, resulting in a dataset of 37 individuals representing six species/subspecies (Fig. 1a, Supplementary Table 1). This allowed us to infer interspecific relationships among the whales. Given that the current estimated population size of Rice’s whale is only ∼50 individuals, our inclusion of 25 Rice’s whale (*B. ricei,* labeled Bric) genomes represents potentially half the population. All *B. ricei* samples were collected in the Gulf of Mexico or the Atlantic Ocean along the southeastern coast of the United States (Fig. 1b). Our dataset also includes two Eden’s whales (*B. e. edeni,* Bede) from the Pacific Ocean, one collected in Australian waters (southern range) and one from Macau (northern range); four Bryde’s whales (*B. e. brydei,* Bbry): two from the Caribbean Sea near the Rice’s whale distribution, one from the Indian Ocean, and one from the eastern North Pacific; a sei whale (*Balaenoptera borealis,* Bbor); a blue whale (*Balaenoptera musculus,* Bmus); and four fin whales (*Balaenoptera physalus*, Bphy) (Fig. 1a).

**Fig. 1.**
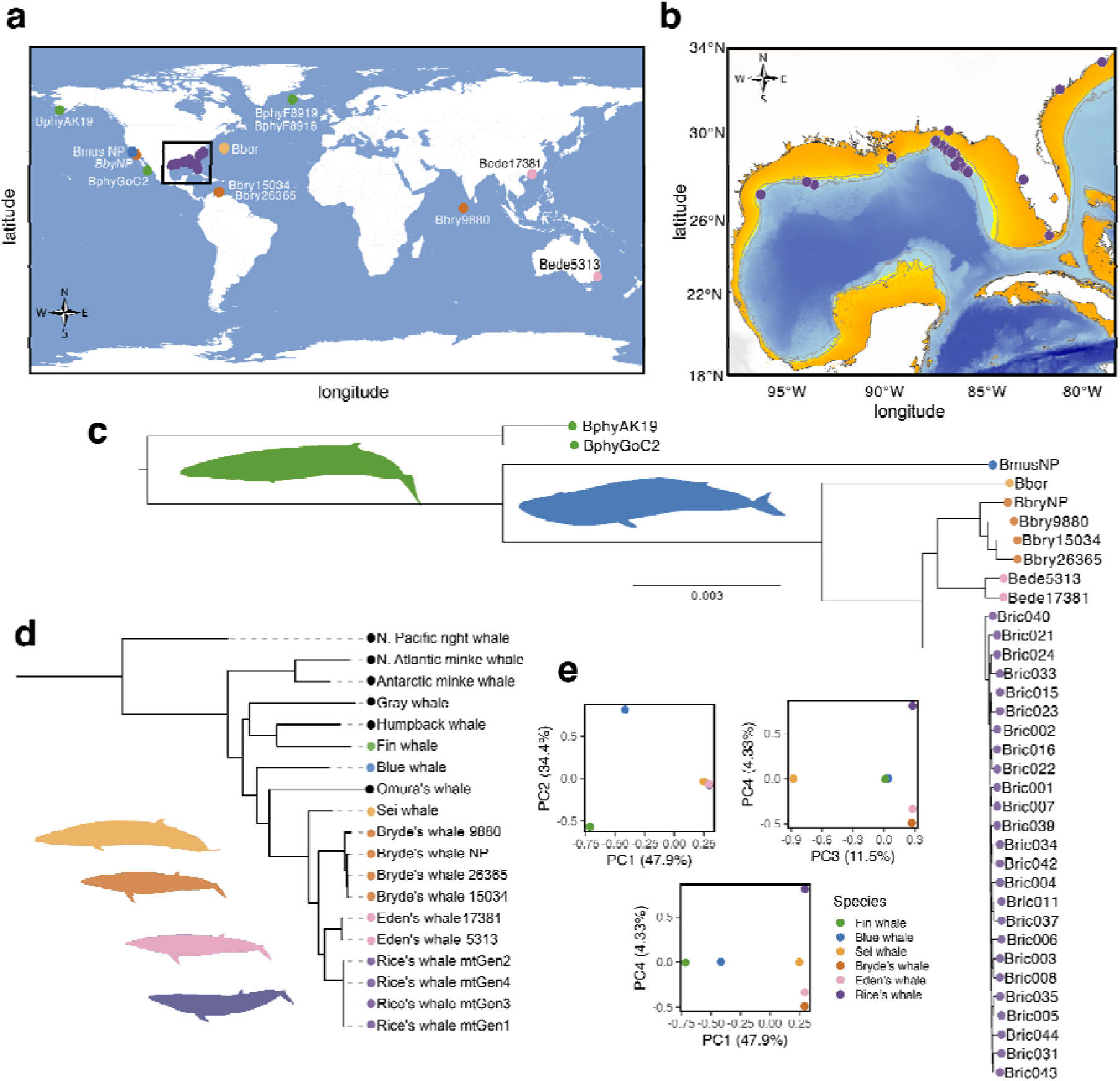
Phylogeny and genetics of baleen whales. **a,** Locations of Rice’s whales (purple dots), Eden’s whale (pink), Bryde’s (dark orange), sei (light orange), fin (green) and blue (blue) whale samples. **b,** Bathymetric map of the Gulf of Mexico showing lines that connect all points of equal depth (isobaths) for -100m and -400m (gray). The transition to yellow is at -140m, indicating the regions of the Gulf of Mexico that were likely above sea level during the last several glacial maxima (∼22,000ybp, 140,000ybp, and several earlier glacial periods through the late Pleistocene). Current primary habitat of Rice’s whales is between -100m and -400m isobaths in the northeastern Gulf of Mexico, extending to some regions of the northwestern and southwestern Gulf of Mexico^5,6^. **c,** Nuclear phylogeny based on 5,039 BUSCO loci. All nodes had 100% bootstrap support except for some within the Rice’s whale clade. **d,** the mitochondrial phylogeny based on complete mitogenomes. All species and subspecies clades had >95% bootstrap support. **e,** PCA using one individual per species.

A phylogeny based on orthologous nuclear genes recovered the same topology as the only other genomic analysis of the Bryde’s whale complex ^16^, with Eden’s and Bryde’s whales as sister taxa and *B. ricei* as the outgroup (Fig. 1c). This topology differs from mitochondrial phylogenies based on the control region ^5^ and whole mitogenomes, which places Eden’s and Rice’s whales as sister taxa, with the Bryde’s whale as the outgroup (Fig. 1d). These results underscore the utility of large nuclear datasets to resolve evolutionary relationships among closely related taxa.

We next investigated population structure using principal components analysis (PCA), subsampling one individual per species (Fig. 1e). The results are consistent with the phylogeny: PC1 (47.9%) separates the fin whale from the sei whale and Bryde’s whale complex (Rice’s, Eden’s and Bryde’s whales), PC2 (34.4%) separates blue whales, PC3 (11.5%) separates the sei whales and PC4 (4.33%) distinguishes the Rice’s whales from Eden’s and Bryde’s whales. ADMIXTURE ^22^ analysis showed a similar pattern, the best-supported model (K = 3) separated Rice’s whales from the rest of the baleen whale species (Supplementary Fig. 1b). At higher K values, members of the Bryde’s whale complex form separate clusters, although sparse sampling for some species limits the resolution ^23,24^. Together, these results support the phylogenetic distinctiveness of the Rice’s whale relative to other baleen whales.

### Genetic diversity and signatures of inbreeding

Runs of homozygosity (ROH) are contiguous stretches in the genome that are homozygous due to inheriting identical haplotypes from both parents in that region ^25^. The fraction of the genome in ROH (F_ROH_) can serve as a measure of genome health ^26^ and long ROH in particular (>>1 Mb) are a reliable indicator of inbreeding depression ^27^. To characterize patterns of genetic diversity in Rice’s whales, we estimated genome-wide heterozygosity and F_ROH_ and compared these metrics across other baleen whale species (Fig. 2a). Rice’s whales show the lowest genome-wide heterozygosity, with an average of 1.34 x10^-4^ (n = 25; range: 6.50 x10^-5^ - 1.90 x10^-4^). Heterozygosity is slightly higher in Eden’s whales (n = 2; 2.83×10□□), followed by Bryde’s whales (n = 4; 5.81 x10^-4^), sei whales (n = 1; 8.15x10^-4^), and an order of magnitude higher in fin whales (n = 4; 1.32 x10^-3^) and blue whales (n = 1; 2.99 x10^-3^). Rice’s whales exhibit substantially higher F_ROH_ than other balaenopterids (Fig. 2a), with every sequenced individual showing 22–40% of their genome in ROH. In contrast, all other species have 0.0-0.78% of their genome in ROH, with the exception of an Eden’s whale individual, which has 12.81% of its genome in ROH. However, Rice’s whale genomes contain few ROH longer than 10 Mb (average FROH_10M_ = 2.14%), indicating little evidence for very recent inbreeding (Fig. 2b, Supplementary Table 2). Most ROH fall between 2–10 Mb, followed by a substantial number of smaller ROH (1–2 Mb). ROH of 2–10 Mb suggest some inbreeding within Rice’s whales due to common ancestry dating back to ∼100 to 500 years ago (5-25 generations), suggesting small population size over the last few centuries.

**Fig. 2.**
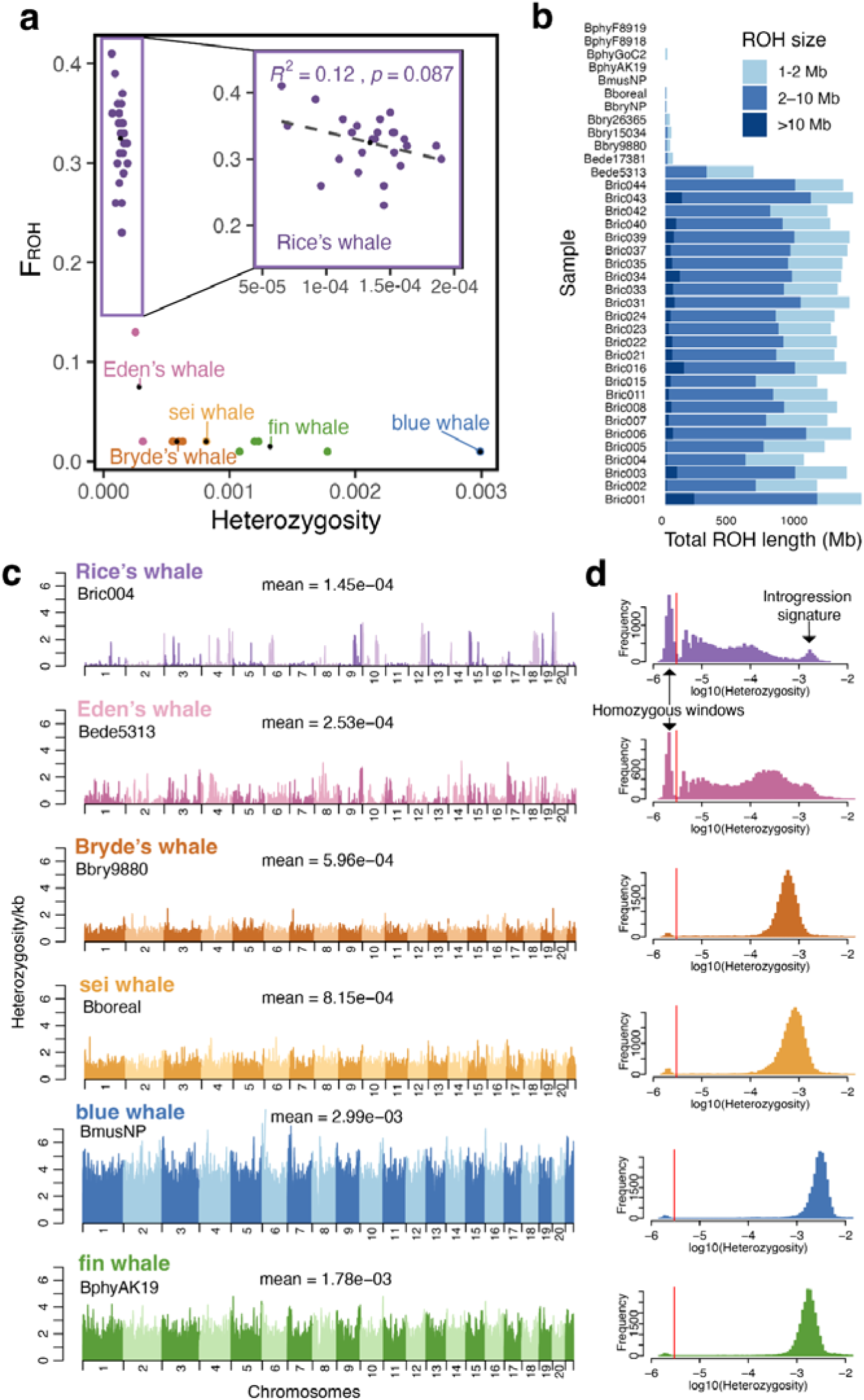
Inbreeding and patterns of genome-wide heterozygosity. **a,** Genome-wide heterozygosity and the fraction of the genome in runs of homozygosity (F_ROH_) per individual. Colors represent the species and correspond to the names of the colored labels. Average per species shown with a black data point. Inset shows a zoom-in of Rice’s whale (*B. ricei*) individuals, with a linear regression line; R² indicates the proportion of heterozygosity variance explained by F_ROH_. **b,** Runs of homozygosity (ROH) size distribution per individual. **c,** Heterozygosity per kb in 1Mb windows from a representative individual from each species. **d,** Logarithmic scale histogram of heterozygosity per kb for 1Mb windows in each individual shown in **c**. The red line denotes cutoff for calling runs of homozygosity.

### Signatures of introgression in spatial heterozygosity

We examined spatial patterns of heterozygosity along the genome (Fig. 2c) by calculating heterozygosity in 1□Mb windows. Most baleen whale species exhibit relatively even heterozygosity, without pronounced runs of homozygosity or peaks of elevated heterozygosity (Fig. 2c), consistent with the ROH statistics (Figs. 2a and 2b). In blue whales, fin whales, sei whales, and Bryde’s whales, heterozygosity distributions show a unimodal, approximately normal distribution, with few homozygous windows (Fig. 2d). In contrast, Rice’s and Eden’s whales display an enrichment of regions with very low heterozygosity, requiring a logarithmic scale to visualize the distribution effectively. Examining the spatial pattern across the Rice’s whale genome reveals some localized peaks of higher heterozygosity (>1×10□³), interspersed with many windows of low heterozygosity (∼5 x10^-5^). Such windows of low heterozygosity are a signature of small, isolated populations.

Particularly in Rice’s whales, we observed distinct peaks in heterozygosity that are clearly elevated above the baseline (Fig. 2c). Although the intensity varies, all Rice’s whale genomes displayed this pattern (Supplementary Fig. 2), characterized by sparse, distinct regions of higher heterozygosity comprising 4.5% of the genome on average among individuals. Initially, we considered that these peaks could reflect artifacts in the reference genome. However, because all outgroup species were also mapped to the same reference and the peaks are not shared across all Rice’s whale individuals, this explanation is unlikely. Instead, these regions could represent introgression from a deeply diverged source ^28,29^, which, upon interbreeding with Rice’s whale, would produce localized regions of elevated heterozygosity.

### Heterozygosity peaks are explained by historical admixture

Our analyses of genome-wide diversity in Rice’s whales suggest a complex demographic history involving both long-term small population size in isolation and recent introgression. To investigate this history, we performed demographic inference using both Approximate Bayesian Computation (ABC) and site-frequency-spectrum-based inference in ∂a∂i. We found that models involving only population size changes provided substantially poorer fit to the observed genomic data than models incorporating admixture from an as-yet unidentified sister “ghost” lineage (Fig. 3a). Both inference methods suggest that the best-supported model consists of a three-epoch demographic history with long-term small effective population size and a recent limited pulse of introgression from a ghost population.

**Fig 3.**
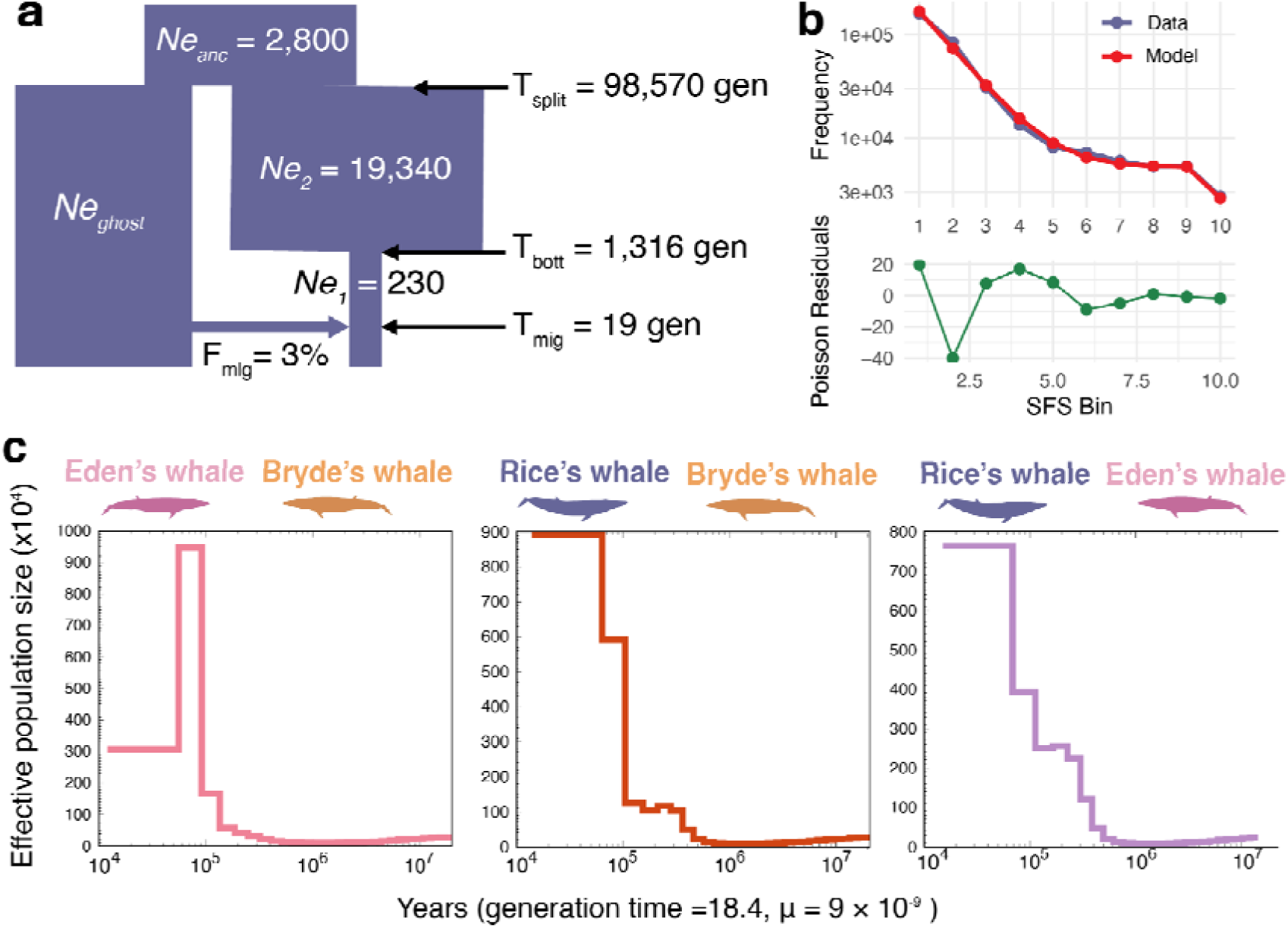
Demographic modeling. **a,** Best model parameters based on ABC and ∂a∂i inference methods. **b,** Top panel shows the folded SFS of the data and the best model from ∂a∂i, bottom panel shows poisson residuals between model and data. **c,** Hybrid pairwise sequential pairwise coalescent (hPSMC), cessation of gene flow between diverging taxa is indicated by an exponential increase in effective population size from the pseudo-hybrid genome (e.g., Eden’s/Bryde’s). Episodic or ongoing low levels of gene flow after the onset of divergence are expected to result in a more gradual or stepped increase in the inferred population size (e.g., Rice’s/Bryde’s, Rice’s/Eden’s). The mutation rate (µ) is in substitutions/site/generation.

ABC posterior estimates (Supplementary Fig. 3, Supplementary Table 3) indicate that the contemporary effective population size of Rice’s whales has remained very small for 2,018 generations (95% credible interval (CI): 985-4,755 generations), with a recent effective population size estimate of *N*_e_ = 236 (95% CI: 175-445). The inferred admixture event was also well constrained, with an estimated admixture proportion of 3.2% (95% CI: 1.8-4.8%) occurring 26 generations ago (95% CI: 18-55 generations). For an effective population size of 236 individuals, this admixture fraction amounts to ∼7-8 effective migrants. Assuming a generation time of 18.4 years, this introgression occurred 331-1,012 years ago. In contrast, deeper demographic parameters were less precisely resolved, including the divergence time between Rice’s whales and the ghost lineage (T_split_ = 47,343 generations; 95% CI: 6,000–104,000) and the effective population size of their common ancestor (*N*_eanc_ = 27,332, 95% CI: 2,000-51,025), which showed broad posterior distributions and strong covariance (*R^2^*= 0.857). Posterior distributions for *N*_eghost_ and the effective population size of Rice’s whales prior to their recent small population size (*N*_e2_) were nearly indistinguishable from their prior distributions, indicating that this approach provided no information about these two parameters.

We next performed demographic inference using ∂a∂i ^30^ under a constrained version of the best-supported ABC model (F_mig_ = 0.03, *N*_eghost_ = *N*_eanc_) to assess concordance between inference methods and further refine our demographic parameter estimates. Under these constraints, ∂a∂i converged on a model (Figs. 3a and 3b) with an ancestral effective population size of 2,800, a divergence time of 98,570 generations (∼1.81 Ma), a pre-bottleneck effective population size of *N*_e2_ = 19,340 and a contemporary effective population size of *N*_e1_ = 230 for Rice’s whales for the past 1,316 generations (∼24.2 ka). The model also timed the admixture pulse to 19.19 generations ago (∼353 years ago; ∼7 effective migrants). Notably, ∂a∂i recovered estimates for *N*_e1_, the admixture fraction, and admixture timing that were highly similar to those obtained from ABC, while also providing estimates for *N*_e2_ and the timing of the bottleneck, which were weakly or not resolved by ABC inference. Thus, ∂a∂i and ABC methods both strongly support a demographic history involving long-term small effective population size in Rice’s whales together with a small amount of historical introgression from a divergent sister lineage.

### Exploring potential sources of introgression

To identify the most likely donor lineage for the observed introgression signal in Rice’s whales, we applied hybrid pairwise sequential Markovian coalescent (hPSMC) ^31^, which models divergence and gene flow between pairs of individuals from different species (Fig. 3c). hPSMC plots show a strong pattern of post-divergence gene flow between both Bryde’s and Eden’s whales with Rice’s whale, and to a lesser extent between Bryde’s and Eden’s whales. It appears that gene flow may have continued among all three species until about 400,000 years ago, but subsequent low-level gene flow or sporadic introgression continued between Rice’s whale and one or both of the other species until 70,000 - 100,000 years ago. While timing estimates from hPSMC carry uncertainty due to sensitivity to mutation rate, generation time, and population structure assumptions, the results are consistent with our demographic inference and help narrow the window of contact between Rice’s whales and their closest relatives.

ABBA-BABA tests using Dsuite ^32^ revealed a significant excess of shared derived alleles between Rice’s whales and Eden’s whales in all individuals (D= 0.12–0.18; p-value < 6.66×10⁻^10^; Supplementary Table 4), using the nuclear phylogeny ((Bryde’s whale, Eden’s whale), Rice’s whale), Fin whale) with fin whale as outgroup (Figs. 1c and 4a). This genome-wide excess of shared derived alleles with Eden’s whales likely reflects shared ancestral variation or ancient admixture with an Eden’s-like lineage; we return to this signal below using branch-specific tests. Comparing the number of ABBA patterns to whole-genome heterozygosity revealed no correlation (Fig. 4b) suggesting that the genome-wide signal is not driving the elevated heterozygosity we observed in Rice’s whales. In contrast, the number of BABA patterns, reflecting shared derived alleles with Bryde’s whales, correlated significantly with whole-genome heterozygosity (R² = 0.22, p-value = 0.019; Fig. 4c), implicating a Bryde’s whale-like lineage as a more likely source of introgression, potentially more recent or localized.

**Fig. 4.**
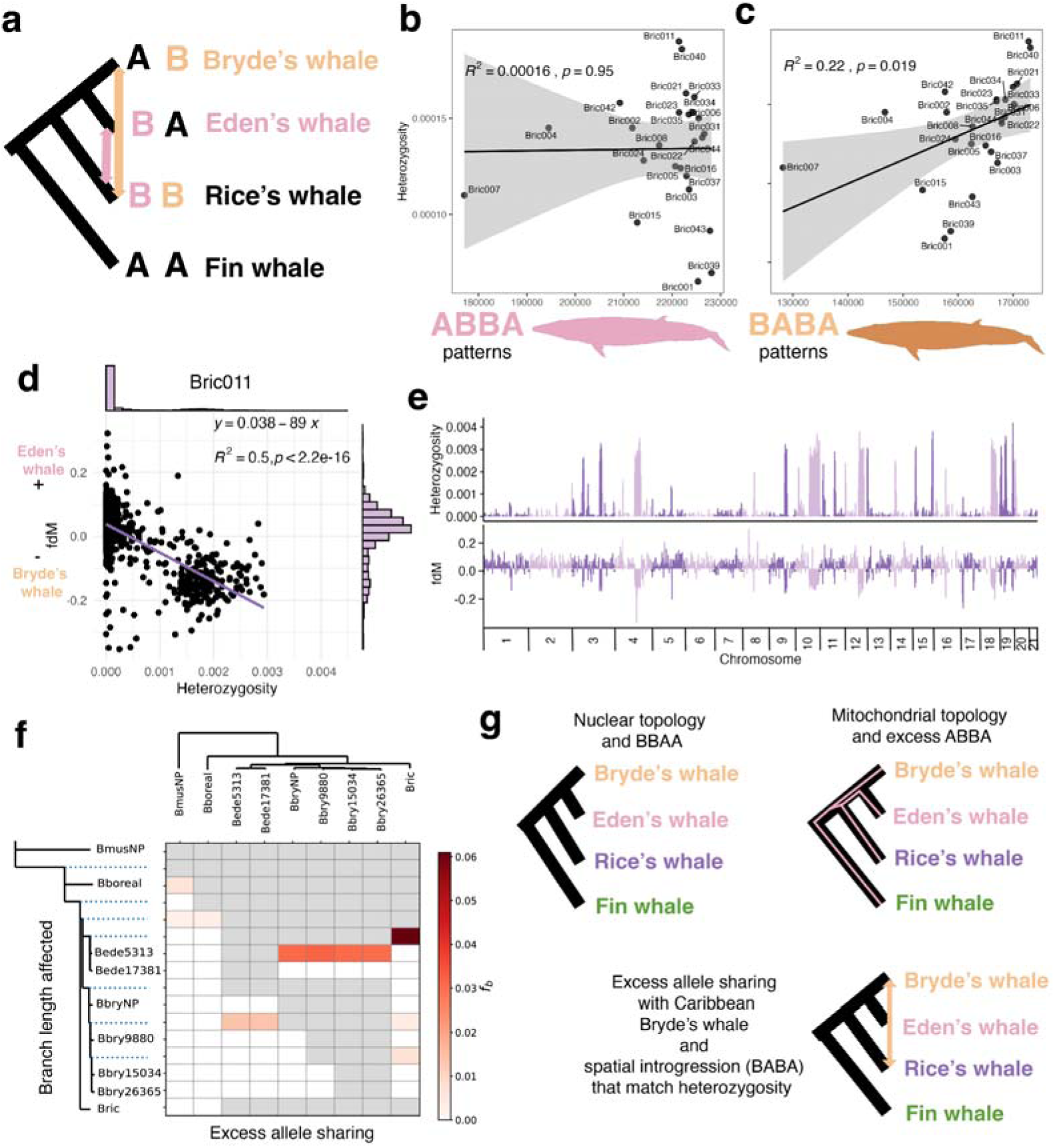
Genomic signatures of allele sharing and introgression in Rice’s whales. **a,** ABBA–BABA tests quantifying excess allele sharing between Rice’s and Bryde’s whales (BABA) and between Rice’s and Eden’s whales (ABBA), using fin whales as the outgroup. **b,** Relationship between the number of ABBA patterns and genome-wide heterozygosity per individual. **c,** Relationship between the number of BABA patterns and genome-wide heterozygosity per individual. **d,** Genome-wide window-based comparison of heterozygosity and fdM in individual Bric011, shown as a scatterplot with marginal histograms; negative fdM values indicate excess allele sharing with Bryde’s whales. **e,** Miami plot of heterozygosity and fdM across the genome of Bric011. **f,** f-branch statistic heatmap showing excess allele sharing across branches of the baleen whale phylogeny. **g,** Schematic trees illustrating the nuclear topology (BBAA), mitochondrial topology (excess ABBA), and introgression, inferred from the spatial concordance of elevated heterozygosity and excess BABA patterns.

To investigate this relationship at finer genomic scales, we calculated *f*dM, in 1Mb windows along the chromosomes, a statistic designed to detect introgression signals locally ^32^. Positive *f*dM values reflect an excess of ABBA patterns, whereas negative values indicate an excess of BABA patterns. Strikingly, all individuals show significant correlation between *f*dM and heterozygosity (Supplementary Table 5), suggesting that introgression is responsible for elevated heterozygosity peaks. For example, Bric011 (r = -0.7, R^2^ = 0.5, p-value < 2.2×10⁻^16^) shows that most windows cluster around both low heterozygosity and fdM values near zero, indicating no introgression signal across the majority of the genome (Figs. 4d and 4e). However a subset of windows in Bric011 (6.88%) exhibit both elevated heterozygosity and negative *f*dM values, consistent with localized introgression from Bryde’s whales. Similar patterns are observed across other Rice’s whale individuals, with the proportion of putatively introgressed windows ranging from 1.44% to 6.88%. Together, these results indicate that the heterozygosity peaks characteristic of Rice’s whales are best explained by localized introgression from Bryde’s whales or a closely related ghost lineage.

To further distinguish ancient shared ancestry from more recent gene flow, we used the *f*-branch method in Dsuite ^32^. Using the nuclear phylogeny (Fig. 1c) and collapsing all Rice’s whale individuals into a single branch, we identified excess allele sharing between Rice’s whales and the ancestral branch leading to Eden’s whales (Fig. 4f), consistent with the genome-wide ABBA excess identified earlier, and likely reflecting ancestral structure in the short internal branch separating Eden’s and Bryde’s whales from Rice’s whales, rather than direct gene flow with Eden’s whales. In contrast, we detected excess allele sharing with Bryde’s whales, particularly with individuals from the Caribbean, suggesting more recent gene flow consistent with the ∼350 year estimate from our demographic modeling. Taken together, these results indicate that both ancient shared ancestry and more recent gene flow, particularly from Bryde’s whales, contributed to the heterozygosity patterns observed in Rice’s whales (Fig. 4e and 4g).

### Introgression did not reduce additive load

One of the challenges faced by small populations is the reduced efficiency of purifying selection due to genetic drift ^33^. In Rice’s whales, fixed deleterious variants were more numerous than in closely related baleen whales, but when looking at the total number of deleterious alleles per individual this difference is attenuated (Fig. 5a, Supplementary Fig. 4). Loss-of-function (LoF) mutations showed more fixed variants in Rice’s whales, but the total number of derived LoF alleles was very similar across species. Normalizing by synonymous mutations revealed a lower proportion of LoF variants (Fig. 5b, Supplementary Fig 4), suggesting that while drift in the small Rice’s whale population may have contributed to the fixation of some deleterious alleles, strongly deleterious mutations were still effectively purged.

**Fig. 5.**
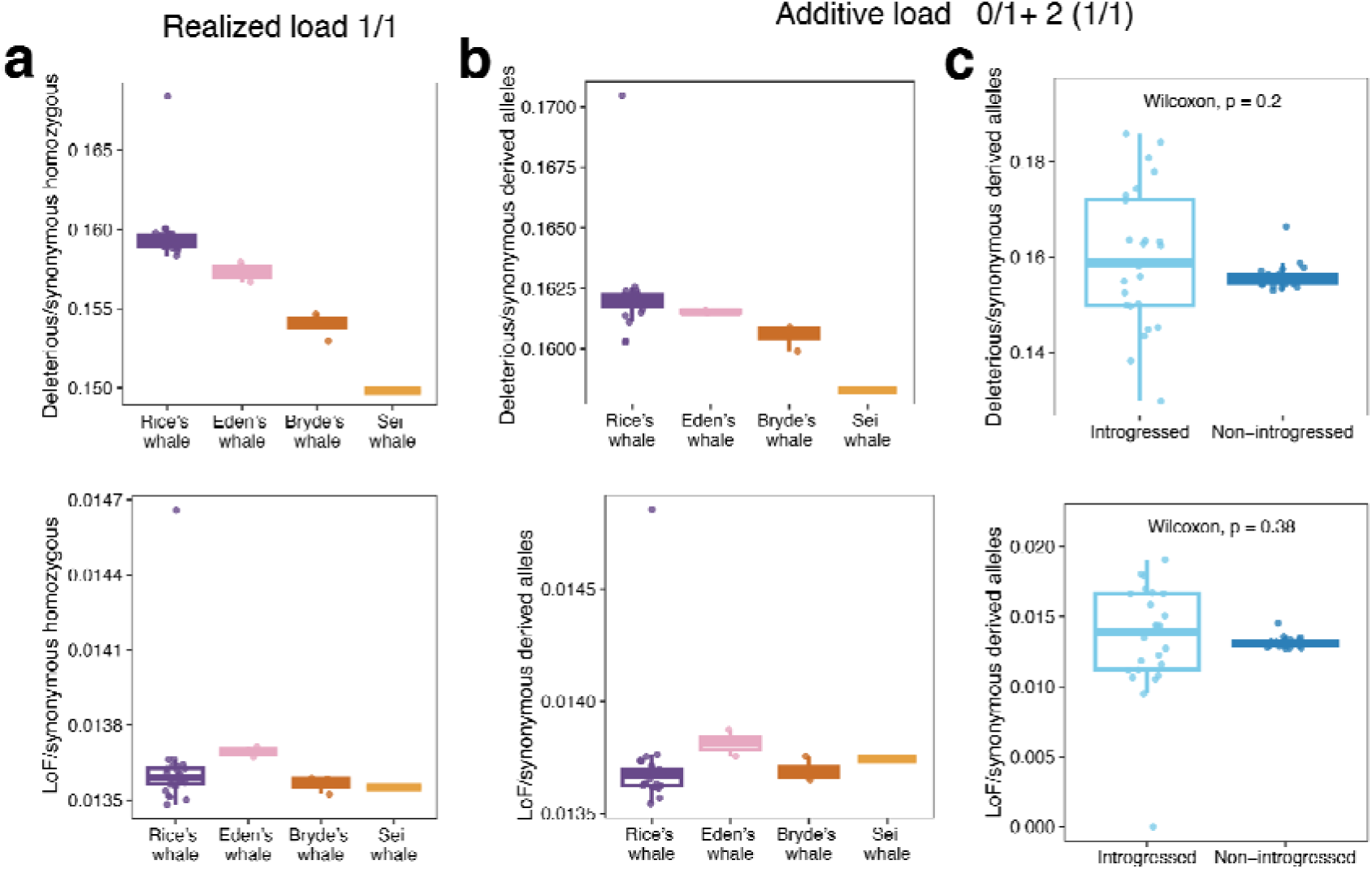
Deleterious and loss-of-function variants in baleen whales. Ratios of deleterious-to-synonymous mutations (top panels) and loss-of-function (LoF)-to-synonymous mutations (bottom panels) are shown for three variant load categories. **a,** Realized genetic load, quantified as the ratio of homozygous derived deleterious or LoF variants to synonymous variants, in Rice’s, Eden’s, Bryde’s, and sei whales. **b,** Additive genetic load, calculated as the sum of all derived deleterious or LoF alleles relative to synonymous variants, across the same four species. **c,** Additive genetic load partitioned by genomic context in Rice’s whales, comparing introgressed regions to non-introgressed regions.

Given the evidence of introgression, we asked whether it might have shaped genome-wide patterns of genetic load ^34^. The total number of deleterious and LoF alleles in the introgressed regions was much lower than what is present in the Rice’s whale genome, which is expected given that the introgressed fraction is small (average 4.5%). The proportion of deleterious and LoF alleles in introgressed regions did not differ from non-introgressed regions, indicating that introgression has not substantially affected the additive load. Introgressed segments have proportionally lower heterozygosity for deleterious and LoF genotypes. However, because introgressed segments are a small amount of the genome, they do not appreciably affect the overall genome-wide load (Fig. 5c, Supplementary Fig 5).

### Modelling the impacts of complex demography on deleterious genetic variation

To further investigate the impacts of long-term small population size, historical gene flow, and recent population declines on genetic diversity and deleterious genetic load in Rice’s whale, we used a forward-in-time simulation model parameterized on our best-fit ABC/∂a∂i demographic model (Fig. 3). We find that prior to the historical migration event 19 generations ago, the ancestral Rice’s whale population had remarkably low genetic diversity (average heterozygosity = 4.5×10⁻□) and relatively high levels of inbreeding and genetic load. The admixture pulse substantially increases heterozygosity to 1.57×10⁻□, and by present day, predicted heterozygosity closely aligns with the empirically observed levels (1.37×10⁻□ vs 1.34×10⁻□; Fig. 6a). Despite these impacts on heterozygosity, we surprisingly observe little or no impact on F_ROH_, genetic load, or inbreeding load (Fig. 6a). These quantities are essentially unchanged following the simulated migration event, contrasting the expectation that migration into small populations is generally expected to alleviate inbreeding and genetic load ^35–37^. However, when modelling a further decline of the Rice’s whale to *N*_e_=25, more substantial genomic impacts are apparent, including significant declines in heterozygosity and increases in inbreeding and genetic load. For instance, within 10 generations at *N*_e_=25, F_ROH_ is predicted to increase from 0.15 to 0.26, resulting in an associated increase in genetic load (Fig. 6a).

**Fig 6.**
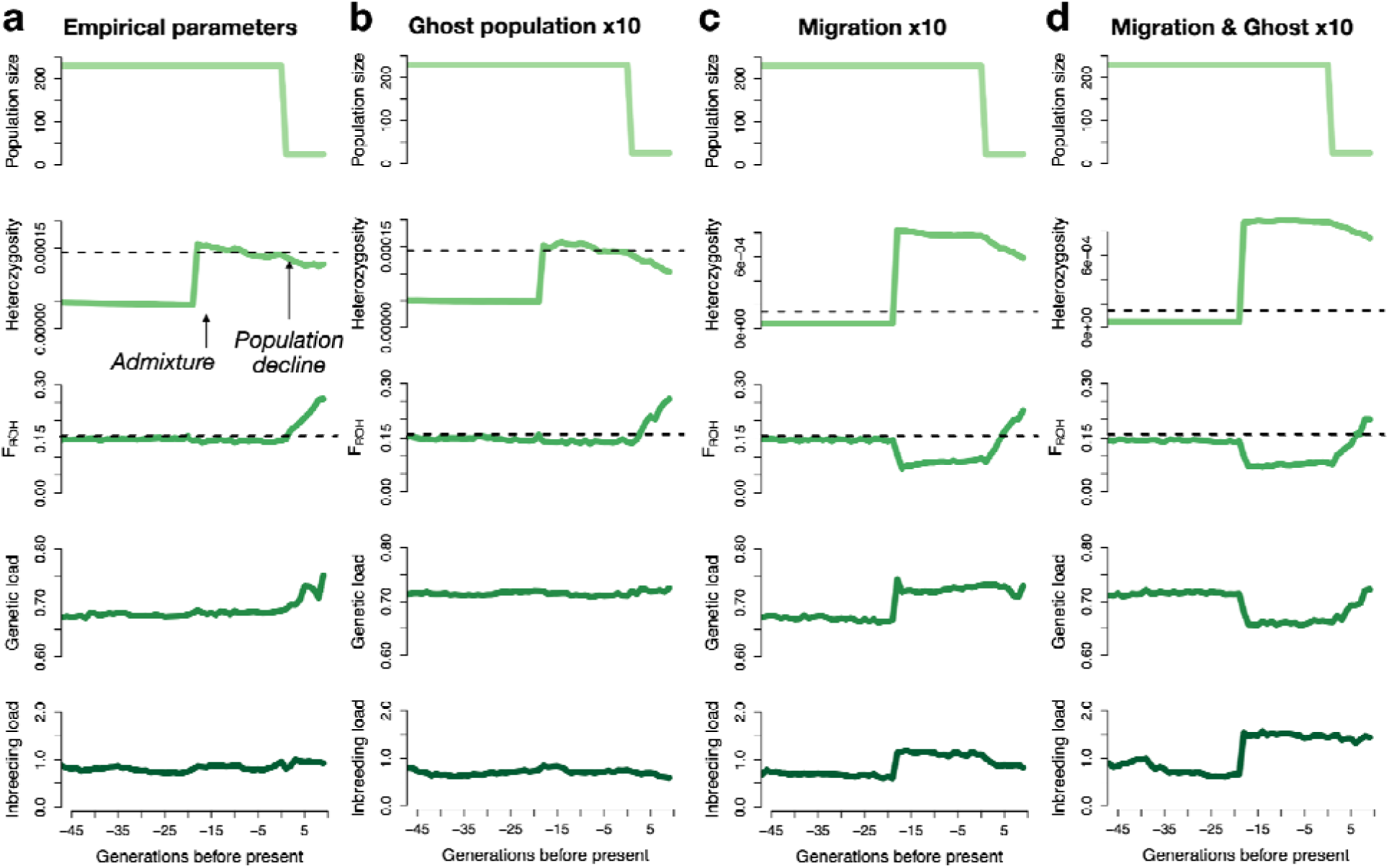
Simulation results highlighting the impacts of complex demography on genomic variation and genetic load. **a**, Results assuming empirically-inferred demographic parameters for the ghost population size and migration rate, **b**, results when increasing the ghost population size 10x to N_ghost_=28,000, **c**, results when increasing the migration rate 10x to F_mig_=0.3, and **d**, results when increasing both the ghost population size and migration rate. For each simulated scenario, results are shown for 45 generations before present and projected 10 generations into the future assuming a hypothetical bottleneck to N_e_=25. Top panel shows effective population size over time for the Rice’s whale population, second panel shows average observed heterozygosity, third panel shows average F_ROH_ for ROH>1Mb, fourth panel shows realized genetic load, and bottom panel shows inbreeding load calculated as the diploid number of lethal equivalents. Dashed lines indicate observed heterozygosity and F_ROH_ for the present day. Note the differing y axis scales for heterozygosity results from panels A and B vs panels C and D. See Supplementary Fig. 6 for results for the ghost population.

We hypothesized that the minimal impacts of admixture on inbreeding and genetic load could be due to both the very low historical migration rate (F_mig_=0.03) and/or relatively small ghost population size (*N*_ghost_=2,800). To explore the impact of these two factors on our model results, we ran simulations with 10x increased ghost population sizes (*N*_ghost_=28,000) and migration rates (F_mig_=0.3). Intriguingly, we observe almost identical results when modelling an increased ghost population size (Fig. 6b), suggesting that the small ghost population size alone does not explain our observations. When increasing the migration fraction 10x, we observe far more substantial genomic impacts, including heterozygosity increases to 7.7×10⁻□, F_ROH_ decreases from 0.15 to 0.10, and notable changes in genetic load and inbreeding load (Fig. 6c).

Importantly, despite these changes, genetic load is observed to increase rather than decrease following admixture. This is explained by the demographic history of the ghost population itself: centuries of small population size had allowed deleterious variation to accumulate in the ghost lineage (Supplementary Fig. 6), meaning that introgressing haplotypes carried their own genetic burden into the Rice’s whale genome. Only when increasing *both* the migration fraction *and* ghost population size do we observe results more typical of a ‘genetic rescue’ (Fig. 6d), whereby heterozygosity increases and F_ROH_ and genetic load decrease, at the expense of an elevated inbreeding load due to an introduction of recessive deleterious alleles ^10^.

Overall, our simulation analysis highlights the often subtle ways in which complex demography can shape genomic variation and deleterious genetic load. We show that, although a very small historical migration pulse can substantially impact genetic diversity, other aspects of genomic inbreeding and genetic load may be less affected. However, regardless of the modelled scenario, we find that ongoing population declines are predicted to result in substantial genomic erosion, characterized by losses of heterozygosity and increases in inbreeding and genetic load. Thus, these findings further reinforce the need to maintain Rice’s whale population sizes to avert irreversible genomic impacts that would compromise the long-term viability of this critically endangered species.

## Discussion

The Rice’s whale presents a unique and instructive case study at the intersection of evolutionary genomics and conservation biology. Our whole-genome analyses reveal a species that has persisted as an exceptionally small, isolated population in the Gulf of Mexico for tens of thousands of years, has experienced a discrete pulse of introgression from a Bryde’s whale lineage ∼350 years ago that reshaped its genetic diversity without meaningfully altering its genetic load, and now faces imminent extinction risk from ongoing anthropogenic pressures. Together, these findings challenge simplistic frameworks for thinking about genetic distinctiveness, genetic rescue, and conservation management, and underscore the importance of integrating complex demographic history into genomic assessments of endangered species.

Species delimitation has long been contentious, and accumulating genomic evidence across cetaceans indicates that introgression is far more pervasive than previously appreciated ^38–41^. Rather than representing rare exceptions, gene flow and reticulate evolutionary histories appear to be common features of whale phylogeny, challenging the notion of genetically “pure” lineages being either typical in natural populations or attainable goals for conservation management. This has important implications for conservation, where management decisions, particularly translocations, are often implicitly guided by assumptions about maintaining genetic integrity, often without explicitly considering the evolutionary ubiquity of gene flow. It is increasingly recognized that admixture is a natural component of evolutionary processes and can play a functional role in shaping genetic diversity ^42–45^.

Our results align with a growing body of work demonstrating that baleen whale evolution is more accurately represented as a network characterized by episodic gene flow between closely related species, rather than as a strictly bifurcating tree ^38–41^. Both the nuclear and mitochondrial phylogenetic trees reflect a fairly rapid radiation of the Bryde’s whale complex, a situation in which mito-nuclear discordance driven by incomplete lineage sorting (ILS) is expected given the short internal branches separating these lineages ^46,47^. Our phylogenetic analysis of nuclear genomic data reveals three well-supported, monophyletic lineages representing Rice’s, Eden’s and Bryde’s whales, with Rice’s whale as the earliest-diverging lineage sister to the Eden’s + Bryde’s clade, consistent with the nuclear phylogeny of Lin et al^16^.

Our demographic inference results suggest that the Rice’s whale has existed as a single small population in isolation in the Gulf of Mexico for tens of thousands of years. Across individuals sampled throughout the northern Gulf of Mexico we find genomic evidence consistent with a single, panmictic population. However, sampling from the western Gulf remains limited (n = 3), and no genomic samples are yet available from Mexican waters where the species has been acoustically detected ^48^. Future sampling should prioritize samples from the southern and western Gulf of Mexico.

Whole-genome analyses reveal a discrete pulse of admixture from a Bryde’s whale lineage ∼350 years ago, contributing ∼4.5% of the genome across sampled individuals. This signal is both discrete and present within all sampled Rice’s whale genomes, consistent with a historical introgression event rather than ongoing hybridization. The majority of the genome therefore reflects a distinct demographic and evolutionary trajectory, with Rice’s whales being more genetically differentiated from Bryde’s and Eden’s whales than the latter are from each other.

Beyond cetaceans, a growing body of work in conservation genomics shows that admixture can be beneficial in small, inbred populations, particularly through its role in increasing heterozygosity and mitigating inbreeding depression. The genetic rescue literature has demonstrated that gene flow can restore fitness by masking deleterious recessive variation, although outcomes are highly context-dependent ^49–51^. In this framework, our study system occupies a particularly informative region of parameter space: although admixture increased the Rice’s whale heterozygosity, we observe neither a reduction nor an increase in its genetic load, contrasting with previously documented cases of genetic rescue in other taxa ^37^. Our simulation results indicate that this outcome is largely governed by the low level of gene flow into Rice’s whale, which mediates the balance between introducing new variation and shifting the frequency of deleterious alleles. At the levels of gene flow inferred here, admixture appears sufficient to elevate heterozygosity but insufficient to meaningfully perturb the distribution of genetic load. This decoupling highlights that increases in genetic diversity do not necessarily translate into immediate fitness gains, nor do they inevitably incur genetic costs. More broadly, these findings reinforce the need to move beyond binary notions of “pure” versus admixed species or populations, and instead evaluate gene flow within a quantitative framework that explicitly considers migration rate, demographic history, and the genetic architecture of load when predicting conservation outcomes.

Like other long-term small populations, including the critically endangered vaquita ^52^, Rice’s whale exhibits low heterozygosity and low genetic load consistent with a prolonged history of small effective population size, with no evidence of recent extreme inbreeding. While the risk of inbreeding depression is relatively low, the currently small population remains at risk of future inbreeding, increased frequency of deleterious mutations through drift, and demographic stochasticity. Our simulations indicate that sustained population growth will be essential to mitigate the risk of future inbreeding following the severe recent decline to fewer than 100 individuals, driven by cumulative anthropogenic pressures including habitat loss, oil spills, vessel noise and traffic, and industrial activity.

The conservation implications of our findings are therefore immediate. Rice whales are unusual among large whales in having an exceptionally restricted distribution and occupying the heavily industrialized Gulf of Mexico, making it ecologically and evolutionarily irreplaceable. Following its formal recognition as a distinct species, additional protective measures were established within the Gulf to reduce vessel strike risk and other threats; however, some of these measures have been subsequently withdrawn. Given the species’ critically small population size and unique evolutionary history, reinstating and strengthening those protections will be essential to secure its long-term persistence.

## Methods

### Sample collection and sequencing

We sequenced 24 Rice’s whale (*B. ricei*) individuals, using skin biopsy samples (n = 19) and tissue from stranded animals (n = 5, 3 collected from Rice’s whales in the northern Gulf of Mexico and 2 from North Carolina and South Carolina in the western North Atlantic) provided by the NOAA SEFSC MMMGL tissue archive (Supplementary Table 1). The two Atlantic samples were obtained from stranded, ill individuals that were presumed to be outside the species’ typical range ^5,53^. DNA was extracted from these samples using standard proteinase K digestion followed by organic extraction ^54^ or using a Qiagen DNeasy Blood and Tissue kit following the manufacturer’s protocol. The DNA extract from one sample was brown in color and was further purified using MO BIO’s PowerClean DNA Clean Up Kit (Qiagen) to remove potential PCR inhibitors. DNA quality was assessed through gel electrophoresis and quantity measured using a DyNA Quant 200 (GE Lifesciences) or Qubit 4 fluorometer (ThermoFisher Scientific, Waltham, MA, USA). Extracted DNA for 20 of these samples was sent to Azenta Life Sciences (Burlington, MA, USA) where sample quality assessment, DNA library preparation, and sequencing was conducted. Genomic DNA was again quantified using the Qubit 2.0 Fluorometer and 100ng of DNA was used for library preparation for each sample.The NEBNext® UltraTM II DNA Library Prep Kit for Illumina (New England Biolabs, Ipswich, MA, USA) was used for library preparation following the manufacturer’s recommendations. Briefly, the genomic DNA was fragmented by acoustic shearing with a Covaris S220 instrument. Fragmented DNA was cleaned up and end-repaired. Adapters were ligated after adenylation of the 3’-ends followed by enrichment by limited cycle PCR. DNA libraries were quantified using a Qubit 2.0 Fluorometer. The DNA libraries were also quantified by real time PCR (Applied Biosystems, Carlsbad, CA, USA). The sequencing libraries were clustered onto Illumina HiSeq flow cells with one sample per lane for approximately 30x coverage. Raw sequence data (.bcl files) generated were converted into fastq files and de-multiplexed using Illumina bcl2fastq 2.17 software. One mismatch was allowed for index sequence identification.

Our dataset also includes two Eden’s whales (*B. e. edeni*) from the Pacific Ocean, one collected in Australian waters (southern range) and one from Macau (northern range). We sequenced three Bryde’s whales (*B. e. brydei*): two from the Caribbean Sea, near the Rice’s whale distribution, and one from the Indian Ocean. DNA from tissue samples of two Eden’s whales and three Bryde’s whales (Supplementary Table 1) was provided by the NOAA SWFSC DNA and tissue archive. These samples and the remaining four Rice’s whale samples were sequenced at Admera Health (South Plainfield, NJ, USA) with the NovaSeq X Plus platform.

Finally, archived genomic sequences from a Rice’s whale (SRR10331559), a sei whale from the Atlantic Coast of the U.S., *B. borealis* (SRR26062094), a blue whale from the eastern North Atlantic, *B.a musculus* (SRR25748633), a Bryde’s whale from the eastern North Atlantic (SRR27003634) and four fin whales, *B. physalus*, one from the Pacific Ocean, one from the Gulf of California, and two from the North Atlantic (SRR18671251, SRR18671250, SRR23615158 and SRR23615133) were downloaded from NCBI’s Sequence Read Archive.

### Quality control and trimming

Raw paired-end sequencing reads were assessed with FastQC v0.11.9 ^55^ using 20 threads. Reports were summarized with MultiQC ^56^ to generate an overview of data quality. Reads were then trimmed with BBDuk (BBTools v38.79) ^57^ using the following parameters: ktrim=r (3′ adapter trimming), k=23, mink=11, hdist=1, tbo (trim adapters based on pair overlap), qtrim=rl (quality trimming from both ends), trimq=15, maq=20, and minlen=40. Trimmed paired-end reads were subsequently processed with repair.sh (BBTools) to ensure that mate pairs were synchronized, with orphan reads written separately. Following trimming, reads were re-assessed with FastQC and summary reports were generated with MultiQC.

### Read mapping and duplicate removal

Trimmed reads were aligned to the *B. ricei* reference genome (GCA_028023285.1_mBalRic1.hap2_genomic.fna) using BWA-MEM2 v2.2.1^58^ with the -M flag (mark shorter split hits as secondary) and -t 10 threads. Read groups were added during alignment with Picard v2.23.9 ^59^ (AddOrReplaceReadGroups), using the following fields: RGID and RGCN set to four letter code of species, i.e. Bric, RGSM = sample ID, RGPL = illumina, and using the SORT_ORDER=coordinate option. Aligned BAM files were processed to index and remove duplicate reads. First, BAM files were indexed with Samtools v1.11 ^60^. Duplicate reads were then removed with Picard v2.23.9 (MarkDuplicates) using 20 GB of memory (-Xmx20G). Picard was run with REMOVE_DUPLICATES=true to remove duplicate reads, and calculate mapping statistics. The resulting deduplicated BAM files were subsequently indexed with Samtools.

Mitochondrial genomes were assembled by first generating a de novo assembly from one sample (Bric006, SRR10331559) using NovoPlasty (v.4.1.pl ^61^), using the North Atlantic right whale mitogenome (NC_037444) as the seed and reference sequences. The de novo mitogenome sequence was rearranged to start with the tRNA-Phe gene and 40bp added from each end to the opposite end to ensure coverage over the artificial ends of the linearized mitogenome sequence when used as a reference sequence for read mapping. We then mapped reads from the remaining samples to the de novo reference using BWA-MEM2, trimmed off the 40bp padded ends, and aligned the mitogenome sequences using MUSCLE (v.3.8.425 ^62^) as implemented in Geneious Prime (v2019.2.3, Biomatters Ltd).

### Variant calling and filtering

The reference genome (GCA_028023285.1_mBalRic1.hap2_genomic.fna) was divided into 25 Mb genomic intervals using Bedtools v2.30.0 ^63^. Per-sample processing included renaming BAM files, indexing with Samtools, and estimating coverage with GATK ^64^ DepthOfCoverage v4.2.0.0 using a minimum mapping quality of 30 and minimum base quality of 20. Variants were called per sample with GATK HaplotypeCaller in BP_RESOLUTION mode, combined with GATK CombineGVCFs, and jointly genotyped with GATK GenotypeGVCFs using the -all-sites and - stand-call-conf 0 options. Variants were then standardized with GATK LeftAlignAndTrimVariants and repetitive regions identified by RepeatMasker and Tandem Repeats Finder were masked using GATK VariantFiltration. Additional custom filtering modified from a previous study^52^, updated to accommodate newer GATK phasing. At the site level, reference alleles and alternate alleles were required to be one of [A, C, G, T,.]; sites lacking AD (allele depth) or DP (depth) fields were removed; and any site with more than 40% missing genotypes across samples was flagged. To remove sites with extremely high heterozygosity, which could reflect bioinformatics errors, we removed sites with >75% het genotypes in 25 Rice’s whale individuals, INFO field thresholds were also enforced, removing variants with QD < 4.0, FS > 60.0, MQ < 40.0, MQRankSum < –12.5, ReadPosRankSum < –8.0, or SOR > 3.0. At the genotype level, individual depth thresholds were set as one-third and twice the mean coverage per sample.

Heterozygous genotypes were only retained if allele balance (REF/DP) was between 20–80%, while homozygous reference genotypes required ≥90% REF reads and homozygous alternate genotypes required ≤10% REF reads; genotypes failing these thresholds were recoded as missing. Sites with no called genotypes were excluded. The final dataset thus contained only high-confidence SNPs, with missingness up to 40% tolerated in outgroup samples (10 of 34 individuals).

### Phylogeny

We generated a maximum-likelihood species phylogeny based on 5,039 nuclear single-copy orthologous sequences identified with BUSCO ^65^ extracted from the Rice’s whale genomes plus those of five outgroup species (Eden’s whale, Bryde’s whale, sei whale, blue whale, fin whale). Specifically, we first used ANGSD ^66^ to generate consensus sequences from the reference-aligned reads (BAM files) of each individual using the following options: *-minQ 30 -minMapQ 25 -doMajorMinor 1 -doMaf 1 -doCounts 1 -GL 1 -doFasta 2 -doDepth 1*. Conserved single-copy orthologous loci were identified in the Rice’s whale reference genome using BUSCO (v5.3.2 ^67^), and the output list of loci was filtered to generate a bed file of only complete, non-duplicated loci between 200 bp and 15 kb in length (n = 5,182). The bed file was used to extract the locus sequences from each consensus sequence using bedtools (v2.31.1 ^68^) with the option ‘getfasta’. Loci with >1% missing data (N’s) were excluded. Sequences within each FASTA file were aligned with mafft (using *--auto*) and trimmed with trimal (using *-automated1*) before being merged into a single FASTA file using catsequences ^69^ for input into IQ-TREE2 ^70^. The full concatenated sequence contained 31,636,815 total sites (393,647 parsimony-informative sites, 257,548 singleton sites, and 30,985,620 constant sites). We ran both partitioned ^71^ and non-partitioned tree inference using ModelFinder ^72^ and 1,000 bootstrap replicates (UFBoot; ^73^) within IQ-TREE2. The tree topologies yielded by the partitioned and non-partitioned runs were identical, but the partitioned tree had a slightly longer total branch length and obtained higher Akaike and Bayesian information criterion scores. The best partitioning scheme and model was an edge-linked-proportional partition model of 319 partitions with separate substitution models and rates across sites. All nodes of the tree had 100% bootstrap support - except for some within the Rice’s whale clade due to the extremely high identity between Rice’s whale sequences - and placed Bryde’s and Eden’s whales within a clade sister to the Rice’s whales.

The phylogeny based on complete mitochondrial genomes was constructed using W-IQ-TREE ^74^, with the best model (TIM2+I+G4+F) based on Bayesian Information Content (BIC). All species and subspecies clades had >95% bootstrap support. Sample and sequence source information are in Supplementary Table 1.

### Population structure

Population structure was evaluated with PLINK v1.9 ^75^, using the command plink --pca with 1) one individual from each species with 28,924,089 variants (Fig. 1e), and 2) with all individuals and 5% minor allele frequency (--maf 0.05), resulting in 15,895,983 variants for principal component analysis (Supplementary Fig. 1a). The PCA including all individuals yielded qualitatively identical interspecific clustering patterns. In that analysis, Rice’s whales exhibit greater dispersion along minor axes, consistent with more extensive sampling of intraspecific genetic variation rather than elevated interspecific divergence. We also used the maximum-likelihood clustering algorithm implemented in ADMIXTURE ^22^. Analyses were conducted from a filtered VCF file containing high-quality SNPs (28,924,089). Variants were first filtered using VCFtools to remove sites with minor allele frequency (MAF) < 0.02 in order to exclude rare alleles that can inflate clustering noise. To reduce the effects of linkage disequilibrium, SNPs were then physically thinned to retain one variant every 10 kb across the genome. ADMIXTURE was run for K = 2-7 ancestral populations using 10 computational threads, and model support was evaluated using cross-validation.

### Relatedness

Relatedness among individuals was assessed with ngsRelate ^76^. First, we extracted the *B. ricei* samples from the final VCF (37 individuals, Rice’s whales and outgroups) using bcftools v1.12 ^77^, restricted to SNPs with a minor allele frequency (MAF) ≥0.05, yielding 489,458 sites in 25 Rice’s whale individuals. We then ran ngsRelate using genotypes calls (-T GT) from the VCF as input to calculate relatedness (*r_ab_*) which uses a formula that accounts for inbred populations^78^. To ensure that downstream analyses were not biased by close kinship, we applied a procedure to exclude individuals so that no pairwise relatedness exceeded *r_ab_* = 0.2. Specifically, we first identified all pairs with *r_ab_* > 0.2 and removed individuals that appeared in multiple such pairs. For pairs where both individuals were only involved in a single related pair, we excluded one individual at random (Supplementary Table 6). This stepwise procedure minimized the number of samples resulting in a final dataset of 13 minimally related *B. ricei* individuals (Bric002, Bric004, Bric008, Bric015, Bric023, Bric031, Bric033, Bric034, Bric035, Bric039, Bric040, Bric042, Bric044).

### Heterozygosity and runs of homozygosity

We calculated heterozygosity as the proportion of all called genotypes in an individual that are heterozygous. This metric reflects the average number of pairwise differences per site between the two homologous chromosomes of that individual. The denominator encompasses all positions with called genotypes, including those that are invariant and identical to the reference. We calculated per-window heterozygosity from genotype call data. To do this, for each individual, we counted the number of heterozygous sites and total called sites in 1 Mb genomic windows with a 100 kb step. We retained only windows with at least 20% of sites successfully called (i.e., >200,000 called genotypes per 1 Mb window). Heterozygosity for each window was computed as the ratio of heterozygous genotypes to called sites.

To quantify runs of homozygosity (ROH), we used two complementary approaches: *bcftools* roh ^77^ and a custom window-based script. In our custom method, we first defined homozygous windows as those with heterozygosity ≤ 3 x10^-6^. We also evaluated two alternative thresholds (≤ 5 x 10^-6^ and ≤ 150 x 10^-6^) to assess the sensitivity of our results to different definitions of homozygosity. For each individual, the fraction of the genome in ROH (F_ROH_) was calculated as the proportion of windows meeting the homozygosity threshold relative to the total number of analyzed windows.

For *bcftools* roh, we provided only variant sites, set the -G 30 option, and specified that ROH should be inferred using allele frequencies estimated from the unrelated Rice’s whale individuals. To estimate the maximum possible ROH length for normalization, we included a pseudo-genome composed entirely of homozygous genotypes when re-running bcftools roh. This pseudo-genome did not affect ROH calls for the real individuals but allowed us to obtain the maximum total ROH length expected for a fully homozygous genome. The summed ROH length of the pseudo-genome (2,397,273,255 bp, slightly shorter than the full autosomal genome length of 2,422,085,147 bp) was then used as the denominator when calculating F_ROH_. The results from both approaches were highly consistent for Rice’s whales when using the ≤ 5 x10^-6^ threshold and *bcftools* estimates (Supplementary Table 2). However, bcftools was unable to estimate ROH for the outgroup species because the method requires a background heterozygosity estimate derived from unrelated individuals of the same species, which we lacked for the non–Rice’s whale samples. For the main text, we report results based on the ≤ 3 x10^-6^ threshold because examination of the log-transformed heterozygosity distribution (Fig. 3) revealed a clear separation between homozygous and heterozygous windows, supporting this cutoff. When calculated with *bcftools* ^77^, all Rice’s whales have 30–50% of their genome in ROH. If we restrict our definition to windows with zero heterozygous sites, Rice’s whales still show 11–25% F_ROH_ (Supplementary Table 2).

To estimate the age of ROH, we used a model in which ROH length decays exponentially through recombination over time ^79,80^. In this framework, the number of generations to the common ancestor (g) is approximated as *g* = 100/(2 × *L*) ^81^,where *L* is the ROH length in centimorgans (cM).

### Projection of site frequency spectrum

A vcf file comprising only putatively neutral SNPs from unrelated individuals was used to obtain the site frequency spectrum (SFS). We filtered out SNPs that fell within coding regions (+-20kb) and with uneven coverage. We also excluded regions in CpG islands, repetitive regions and conserved regions (compared to *Danio rerio*). To avoid uncertainties in ancestral state classifications, we computed a folded SFS. This SFS was calculated based on a projection of the number of segregating sites in unrelated individuals using easySFS (https://github.com/isaacovercast/easySFS) ^82^. From this projection, an optimal number of haploid individuals with a maximized number of SNPs are identified and this number is then used to construct the folded SFS. In our case we did the projection using ten individuals to maximize data completeness while accounting for missing data. Thereafter, the count of monomorphic sites was calculated and added to the 0-bin.

### Demographic history reconstruction ∂**a**∂**i**

We utilized the projected neutral SFS generated above to reconstruct the demographic history of Rice’s whales using ∂a∂i ^30^ and a similar approach to that used for demographic inference of fin whales ^83^. To explore a variety of possible demographic scenarios, we tested the following models (Supplementary Fig. 3):

1. One-epoch: model with no population size change. This model provides a“null model” that estimates ancestral population size (*N*_eanc_).
2. Two-epoch: model with one size change event, from the ancestral size (*N*_eanc_) to the current size (*N*_e1_) occurring T generations ago.
3. Three-epoch: model with two size change events. The first event changed from the ancestral size (*N*_eanc_), then a second population size (*N*_e2_) and lasted for T_2_ generations. The third and current population size (*N*_e1_) T_1_ generations ago.
4. Three-epoch with admixture pulse: to incorporate introgression informed by our genomic results and the ABC simulations (see below), we included a ghost population, which is an unsampled inferred population that was the source of the detected gene flow. We modeled a single pulse of admixture using the implementation in ∂a∂i (PhiManip.phi_2D_admix_1_into_2), where, *f* represents the admixture (migration) fraction from population 1 to population 2 at the time of the pulse.

The haploid sample size (n = 20) plus 5, 15, and 25 were used as extrapolation grid points. The admixture pulse was implemented using PhiManip.phi_2D_admix_1_into_2, with admixture fraction f fixed to 3% based on independent ABC inference (see below). The ghost population (*N*_eghost_) was set to the same size as the ancestral (*N*_eanc_). Lower and upper bounds of model parameters were imposed based on posterior probabilities of the parameters. We used the optimize_log function as our optimization algorithm, and calculated the multinomial log-likelihood for the expected SFS obtained from each optimization. Best-fit parameter sets of each model were scaled using *N*_eanc_ calculated by the equation θ=4*N*_eanc_μL, where L is the total sequence length of the neutral region (415,663,588 bp), μ is the mutation rate (9e-9 substitutions/generation/bp),and θ is the optimal value of theta for the given model. One hundred replicates of each model were performed with randomized starting parameters to assess convergence of the inferred parameters and composite likelihood.

The one- and two-epoch models generated poor fit between the observed and simulated SFS (LL = −68,236 and −5,134, respectively). Some three-epoch models achieved substantially higher likelihoods (LL = −1,891), but these models were characterized by small ancestral population sizes (*N*_eanc_ = 1,563), followed by unrealistically large population expansions (*N*_e_₂ = 291,447) and a very recent, extreme bottleneck (*N*_e_₁ = 57 at T_bott_ = 8 generations), which we deemed to be biologically implausible. Models incorporating migration were also evaluated given the evidence for introgression. While these models generally fit the observed SFS better than isolation-only models, they involved a large number of parameters. For this reason we implemented the ABC methods described below in order to constrain our models. Parameters that we were unable to resolve using the ABC method include the effective population size of the ghost population (*N*_eghost_), and the population size of the Rice’s whale (Ne_2_) after it split from the ghost population and before it contracted to its most recent size. We ran the three-epoch model with admixture pulse, where we fixed the admixture fraction (3%) and the *N*_eghost_ to be equal to *N*_eanc_ and the solution converged to a model with improved likelihood (LL = −1,317).

To estimate uncertainty around the best-fit parameter estimates, we used the Godambe Information Matrix (GIM) as implemented in dadi (dadi.Godambe.GIM_uncert). We generated 100 bootstrap replicates by dividing the genome into 2 Mb chunks and resampling from those chunks using the per-chromosome VCF files of neutral regions described above. The 95% confidence intervals were computed by using the maximum likelihood parameter estimate (± 1.96 × SE) where SE is the standard deviation returned by the GIM (Supplementary Table 3).

### Approximate Bayesian Computation with coalescent-based genomic simulations

Our initial efforts at resolving the Rice’s whale demographic history using ∂a∂i were challenged by the complexity of the parameter-rich models needed to fit our empirical data and the small size of our dataset (just 13 (mostly) unrelated individuals and a species with very low genetic diversity). We reasoned that incorporating other aspects of Rice’s whale genomic diversity besides the SFS could aid our inference of the species’ demographic history. Specifically, both the amount and the distribution of genetic variation across the genome are highly informative about demographic events: the lengths of runs of homozygosity can be used to infer the timing of inbreeding; the degree of heterozygosity across the genome can be used to infer past population size history; and large peaks of heterozygosity can be used to infer the extent and timing of admixture. We therefore used ABC to leverage multiple aspects of genome-wide variation in Rice’s whales to resolve their complex evolutionary history. In short, our ABC method consisted of simulating genomes under a set of predefined models, calculating summary statistics produced by these models, and then comparing to the values from the sequenced genomes to find models with the closest fit.

For each simulation replicate, we generated a set of 13 Rice’s whale genomes with the same size and genotyping call rate as in our empirical dataset. We used the coalescent simulator MaCS ^84^ to simulate whole genomes with Rice’s whale chromosome lengths and randomized recombination-rate maps derived from the human recombination rate map ^85^. We used a mutation rate of 9e-9 substitutions/generation/bp and an overall recombination rate of 1e-8 crossovers/site/generation. Simulated genotypes were processed to reproduce the call-rate distributions in the sequenced genomes of the 13 least related Rice’s whales, and heterozygosity was quantified in 1-Mb genomic windows across the simulated genomes. For both the empirical and simulated genomes, windows with fewer than 20% callable sites were excluded.

We then evaluated the goodness-of-fit between each simulation and the empirical dataset using multiple heterozygosity-based summary statistics designed to capture both genome-wide diversity levels and the spatial structure of heterozygosity (i.e., peaks of heterozygosity and runs of homozygosity). Specifically, we used the distribution of heterozygosity in 1-Mb windows to estimate the following statistics: 1) the mean genome-wide heterozygosity per individual; 2,3,4) the mean height, width and breadth of “peaks” of heterozygosity (defined as windows with heterozygosity >1e-3); 5) the mean genome-wide heterozygosity per individual *excluding* peaks; 6,7) the mean number and combined lengths of windows with zero heterozygous genotypes; 8) the distribution of the number of heterozygous genotypes per window; 9) the distribution of shared peaks of heterozygosity. For each statistic, the distances between simulated and empirical data were calculated as relative absolute deviations or histogram-based summed deviations. Simulations were ranked across each distance metric, and these ranks were then summed to generate an overall goodness-of-fit score.

With this framework, we first evaluated 8 different demographic models of increasing complexity to perform model selection (Supplementary Table 3). The models were: 1-, 2-, and 3-epoch single-population models; 2- and 3-epoch models incorporating a sister “ghost” population derived from a shared ancestral population; 2- and 3-epoch models including a pulse of admixture from the ghost population during the most recent epoch; and a final 3-epoch ghost-admixture model in which the contemporary effective population size of the Rice’s whale was constrained to be small (<<1,000). We generated 20,000 simulation replicates under each model. Relative model support was assessed by the representation of each demographic model among the 1,000 best-fitting simulations out of the 160,000 total (=8 models * 20,000 replicates). We found that the best-fitting simulations included only those that incorporated admixture from the ghost population: 3-epoch ghost-admixture with a recent population decline (n=908), 3-epoch ghost-admixture (n=51), and 2-epoch ghost-admixture (n=41). We then ran one million simulations of the best-performing model (3-epoch ghost-admixture with a recent population decline) to refine our parameter estimates. Approximate posterior distributions for demographic parameters were obtained from the 1,000 simulations with the lowest composite goodness-of-fit ranks out of the 1 million total (Supplementary Fig. 3). Point estimates were summarized using kernel-density posterior modes, and uncertainty was quantified using equal-tailed 95% credible intervals (Supplementary Table 3). The results of the ABC inference were then used to refine our demographic inference with ∂a∂i by running more constrained models with fewer free parameters.

### hPSMC

Hybrid pairwise sequential Markovian coalescent (hPSMC^31^) was used to investigate divergence time and evidence of ancestral gene flow or introgression. The hPSMC estimates the divergence time from PSMC analysis of pseudo-hybrid diploid genomes constructed by combining haploid consensus genome sequences from pairs of species, derived from WGS reads mapped to the Rice’s whale reference genome. To scale the PSMC results, we used an estimated autosomal mutation rate of 4.90x10^-10^ substitutions/year/bp from the vaquita ^86^, and a generation time of 18.4 years based on the estimate for *B. edeni* ^87^ in the absence of a generation time estimate for *B. ricei*. PSMC-specific parameters were: -p=“4+25*2+4+6”, t=15, - r=5, and -N=25.

### Introgression analysis

We tested for evidence of introgression among Rice’s whales and closely related baleen whale species using Dsuite (v0.5 r57)^32^. Analyses were performed with the Dtrios module, which computes Patterson’s D statistic and related f-statistics across all valid species trios. We used a genome-wide VCF containing biallelic SNPs that passed quality filtering. Species and population assignments were specified in a SETS file, and a rooted species tree (see Phylogeny) was provided to polarize allele patterns. Rice’s whales were treated as individual samples to allow direct comparison of introgression signals among individuals, whereas Bryde’s whales and Eden’s whales were each grouped. Fin whales were used as the outgroup; however, fin whale samples from the Atlantic were excluded due to substantially lower sequencing coverage, which could bias allele frequency estimates and test statistics.

To examine whether genome-wide heterozygosity was associated with variation in introgression signal among Rice’s whale individuals, we tested for correlations between individual heterozygosity and the number of informative ABBA–BABA site patterns. We fit linear models in R to assess the relationship between heterozygosity and counts of ABBA–BABA patterns, and evaluate significance using model summaries and Pearson correlation statistics. Relationships were visualized using scatterplots with fitted linear regressions and 95% confidence intervals, with individuals labeled to facilitate comparison across samples. All statistical analyses and visualizations were performed in R, using the packages ggplot2 ^88^ and ggpubr ^89^.

To localize introgression signals along the genome, we used the Dinvestigate module in Dsuite ^32^. Analyses were performed separately for each Rice’s whale individual using sliding windows defined by numbers of usable SNPs, with windows and step size set to 500 SNPs. Windowed *f*dM estimates were intersected with corresponding heterozygosity windows using BEDTools ^63^ intersect, to enable direct comparison between local introgression signal and local genetic diversity. The resulting data were visualized using scatterplots and histograms, and correlations were assessed using linear models in R. We additionally generated Miami-style plots, with *f*dM values displayed above the genomic axis and heterozygosity below, to compare their genome-wide distributions and spatial concordance.

We performed an f-branch analysis using Dsuite Fbranch ^32^, which summarizes *f*₄-ratio results across all trios and assigns introgression signal to specific branches while filtering out nonsignificant values. The f₄-ratio estimates the proportion of the genome that has been exchanged between lineages by quantifying the fraction of shared derived alleles attributable to gene flow. In this framework, f_b_ represents the branch-specific proportion of the genome showing excess allele sharing beyond that expected from the species tree alone. All Rice’s whale individuals were combined into a single focal lineage, whereas individuals from other lineages were analyzed separately. This design allowed us to test whether introgression signals involving Rice’s whales were more consistent with particular populations. The f-branch statistic was computed using a predefined phylogenetic tree to summarize excess allele sharing along internal branches.

### Deleterious variation

The Rice’s whale, Eden’s whale, Bryde’s whale, and sei whale samples were mapped to the Blue whale reference genome (GCF_009873245.2_mBalMus1.pri.v3) ^90^, a VCF was generated following the same steps that were described above using the Rice’s whale genome. Additionally, the VCF was annotated with SnpEff v5.2 ^91^ (custom *B. musculus* database), and functionally scored with SIFT4G ^92^ (custom database). We used a custom script to extract genomic coordinates annotated by SIFT and SnpEff, including all coding sites (CDS), loss-of-function (LoF), missense, deleterious, tolerated, missense and synonymous variants. When multiple annotations were assigned to a site, we retained the most deleterious category according to the following hierarchy: LoF > deleterious > tolerated > synonymous.

For each individual, we then quantified the number of variants falling into each category and classified genotypes as missing (./.), heterozygous (0/1), homozygous reference (0/0), or homozygous alternative (1/1). To account for variation in coverage and callability across individuals, we normalized each functional category by the number of called CDS sites per individual and scaled by the mean number of called sites across all individuals. Specifically, for each individual *i* and annotation category *a*, we calculated:

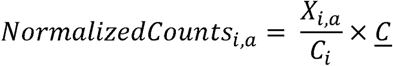

Where *X_i,a_* is the number of variants in category *a*, *C_i_* is the number of called sites in individual *i* and *<u>C</u>* is the mean number of called sites in all individuals. For each category, we quantified: (i) the number of heterozygous genotypes (0/1), (ii) the number of homozygous derived genotypes (1/1), (iii) the total number of derived genotypes (0/1 + 1/1), and (iv) the total number of derived alleles, calculated as:

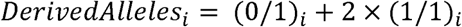

where genotypes were coded relative to the ancestral state (Blue whale reference genome), such that 0 denotes the ancestral allele and 1 denotes the derived allele.

To assess the amount of deleterious variation that falls within introgressed regions, we used the *f*dM statistics per window to determine introgressed regions per individual. We defined as introgressed the windows with values *f*dM <-0.1 and categorized all windows into introgressed/non-introgressed according to this threshold. We converted these coordinates based on the Rice’s whale genome to the Blue whale reference genome using the LiftOver tool from the UCSC Genome Browser ^93^. These regions were intersected with the coordinate files corresponding to the different annotations described above. We normalized the counts using the CDS counts of each individual in each category (introgressed/non-introgressed). Then, we counted the number of homozygotes, heterozygotes, alleles and genotypes as described above.

### Forward simulations

We performed forward-in-time genetic simulations using SLiM 5 ^94^. We simulated a 15 Mb chromosomal segment with randomly generated introns, exons, and intergenic regions following the approach from Mooney et al. ^95^. Exons comprised ∼1% of the total chromosome, thus each segment contained ∼0.15 Mb of coding sequence. We assumed a uniform recombination rate of 1e-8 crossovers per site per generation and a mutation rate of 9e-9 mutations/generation/bp, 70% of which were assumed to be deleterious within coding sequence ^96^ and 4% of which were assumed to deleterious within non-coding regions ^97^.

Selection (*s*) and dominance (*h*) coefficients for nonsynonymous mutations followed the model described in Kyriazis et al ^98^. The majority of new mutations (99.5%) were drawn from a gamma distribution with mean selection coefficient *s* = -0.0131 and shape parameter 0.186, while the remaining 0.5% were fully recessive lethal mutations with *s* = -1.0. Dominance coefficients were modeled as a function of selection strength, with *h* = 0.45 for weakly deleterious mutations (*s* > - 0.001), *h* = 0.2 for moderately deleterious mutations (-0.001 ≥ *s* > -0.01), *h* = 0.05 for strongly deleterious mutations (-0.01 ≥ *s* > -0.1), and *h* = 0.0 for lethal and semi-lethal mutations (*s* ≤ −0.1). This parameterization yields a mean dominance coefficient of *h* = 0.28 for newly arising nonsynonymous mutations, consistent with empirical estimates from human genetic data ^99^. For non-coding deleterious mutations, selection coefficients were drawn from a gamma distribution estimated by Torgerson et al. ^100^ consisting of a mean *s* = −0.001036 and shape = 0.0415. All non-coding deleterious mutations were assumed to be partially recessive (*h* = 0.4).

We assumed demographic parameters for a two population model from our ABC / ∂a∂i inference (Fig. 3), including *Ne*_anc_= 2,800, *Ne*_ghost_= 2,800, *Ne*_2_= 19,340, *Ne*_1_= 230, T_split_= 98,570, T_mig_= 19, T_bott_=1,316, and F_mig_=0.03. F_mig_ determines the migration fraction from the ghost population to N_1_, occurring for a single generation at T_mig_= 19. For each simulation replicate, we ran burn-ins for 10*N_anc_ generations to allow deleterious and neutral mutations to reach equilibrium. Fixed deleterious mutations were discarded during the burn-in but were kept for the remainder of the simulated demography.

With this model, we modified our empirically-inferred demography to explore several hypothetical scenarios. First, to explore the impacts of increasing anthropogenic pressures in the Gulf of Mexico, we modelled an additional bottleneck of the Rice’s whale population to Ne=25 starting in the present day and projected into the future for 10 generations. Second, to explore the impacts of a higher migration rate from the ghost population, we increased the migration fraction 10x from 0.03 to 0.3. Third, to investigate how admixture from a larger ghost population might have impacted Rice’s whale, we increased the ghost population size by 10x from 2,800 to 28,000. Finally, we modelled a combination of the two above scenarios in which both the migration fraction and ghost population size were increased 10x.

We ran 50 replicates for all simulated scenarios. For each replicate, we outputted the following statistics every generation during the final 70 generations of the simulation from both the Rice’s whale population and the ghost population: mean heterozygosity, F_ROH_ for ROH>1Mb from a sample of 80 individuals, mean genetic load (calculated as 1 - multiplicative fitness), and mean inbreeding load (calculated as the diploid number of lethal equivalents ^98,101^). For genetic load and inbreeding load, we projected our results from the 15Mb segment to that of 2.7Gb genome by multiplying the inbreeding load by 180 (as selective effects are summed across sites for inbreeding load) and exponentiating fitness by 180 to compute the genetic load as 1-fitness (as fitness is multiplicative across sites).

## Data availability

The raw sequencing reads generated in this study have been deposited in the NCBI Sequence Read Archive under BioProject accession number PRJNA1246661. Whole-genome VCF data aligned to the chromosome-level assemblies of Rice’s whale (*B. ricei*) and blue whale (*B. musculus*) are available in the Dryad Digital Repository: https://doi.org/10.5061/dryad.s7h44j1q5. All other data supporting the findings of this study are available within the paper and its supplementary information.

## Code availability

All custom code used in this study is publicly available on GitHub at https://github.com/aguilar-gomez/bricei_genomics/

## Acknowledgement

The scientific results and conclusions, as well as any views or opinions expressed herein, are those of the authors and do not necessarily reflect those of NOAA or the Department of Commerce. This research was carried out [in part] under the auspices of the Cooperative Institute for Marine and Atmospheric Studies (CIMAS), a Cooperative Institute of the University of Miami and the National Oceanic and Atmospheric Administration, cooperative agreement #NA25OARX432C0018 and #NA20OAR4320472. This work used computational and storage services associated with the Hoffman2 Shared Cluster provided by UCLA Office of Advanced Research Computing’s Research Technology Group. Portions of this work were performed on the Wynton HPC Co-Op cluster, which is supported by UCSF research faculty and UCSF institutional funds.

## Author contributions

D.A-G., J.A.R., C.C.K., S.N-M., P.E.R., P.A.M., and K.E.L. conceived the study. D.A-G. performed variant calling, heterozygosity, runs of homozygosity, introgression, deleterious variation analyses, demographic inference with dadi, figure generation, and wrote the manuscript. J.A.R. performed ABC simulations and phylogenetic analysis. C.C.K. performed forward-time simulations. M.K. assisted with data analysis. S.N-M. assisted with demographic inference. N.L.V., L.W.T., and P.E.R. provided samples and contributed Rice’s whale expertise to discussions throughout the project. B.Y.K. prepared DNA libraries for sequencing. P.A.M. assisted with data curation, performed hPSMC analysis, and mtDNA analysis. K.E.L. supervised the project. D.A-G., J.A.R., C.C.K., S.N-M., N.L.V., L.W.T., R.D.H., P.E.R., P.A.M., and K.E.L. contributed to the interpretation of results. J.A.R., C.C.K., M.K., P.E.R., and P.A.M. contributed to writing the manuscript. All authors reviewed and commented on the manuscript.

## Supplemental Figures

**Supplemental Fig. 1.**
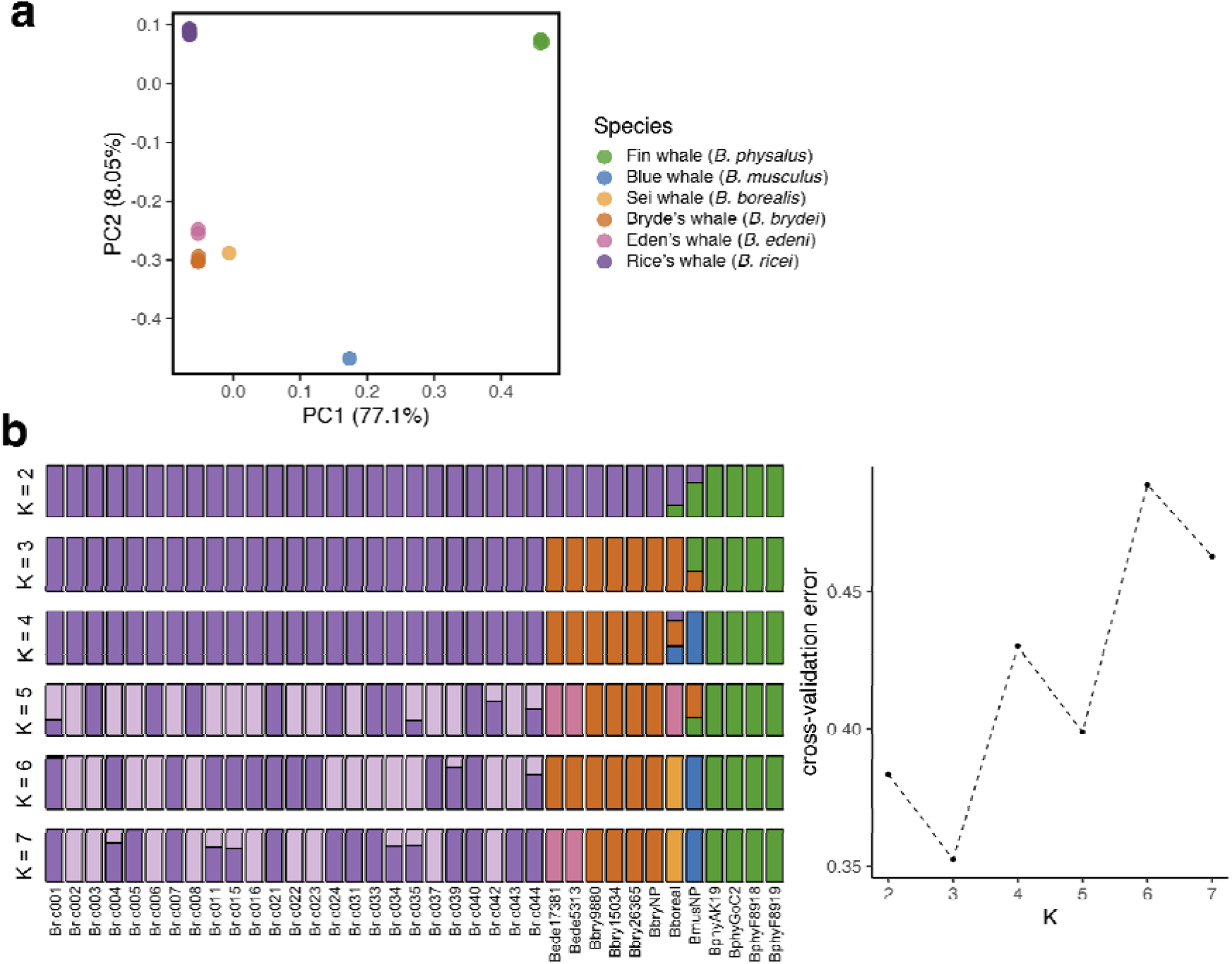
Population structure. **a,** PCA with all individuals, minor allele frequency filter 5% **b,** ADMIXTURE analysis showing k values 2-7 and cross validation analysis

**Supplemental Fig. 2.**
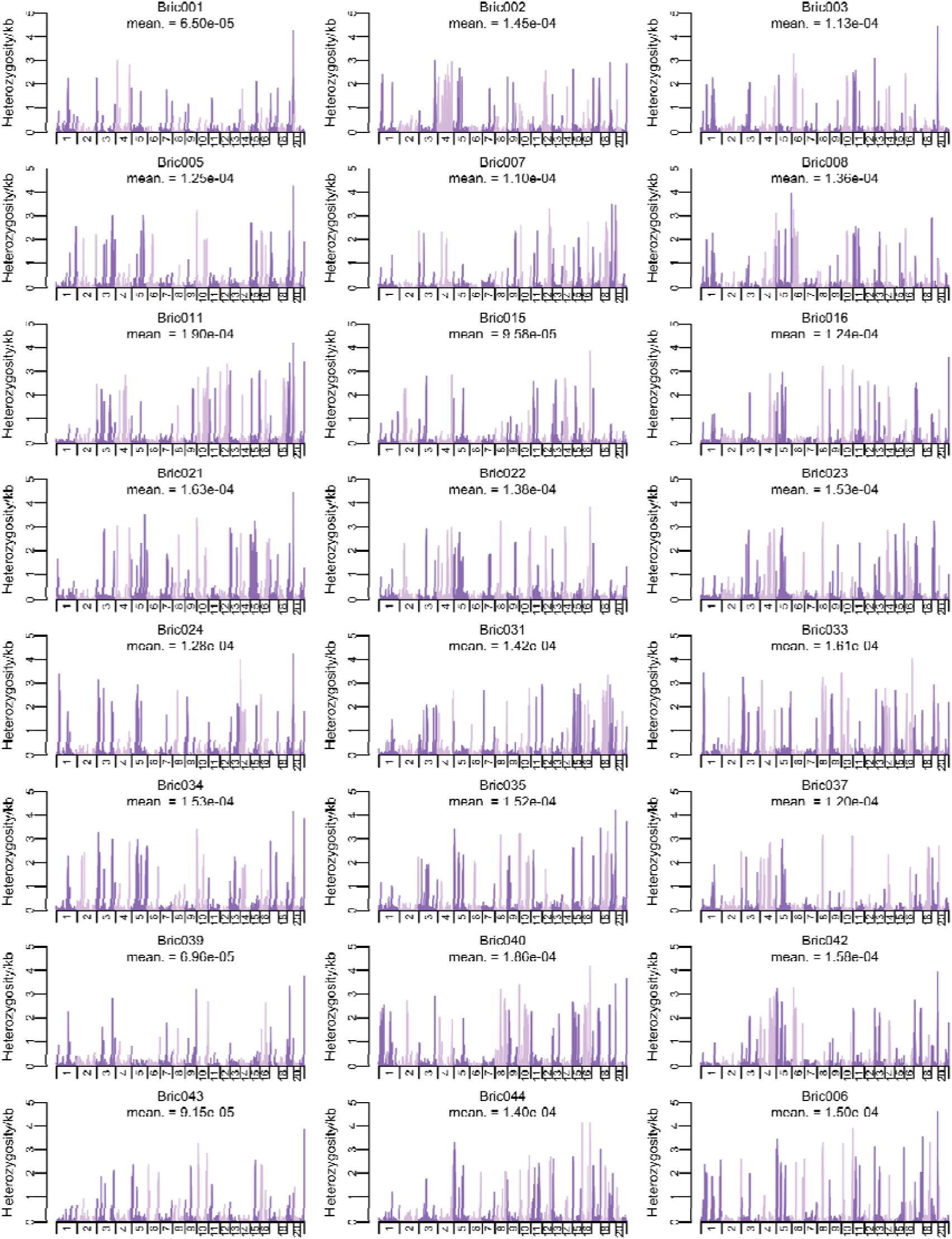
Genome-wide heterozygosity patterns in Rice’s whales. Heterozygosity per kb in 1Mb windows in all sequenced Rice’s whale individuals. Individual Bric004 is in the main Fig. 2

**Supplemental Fig 3.**
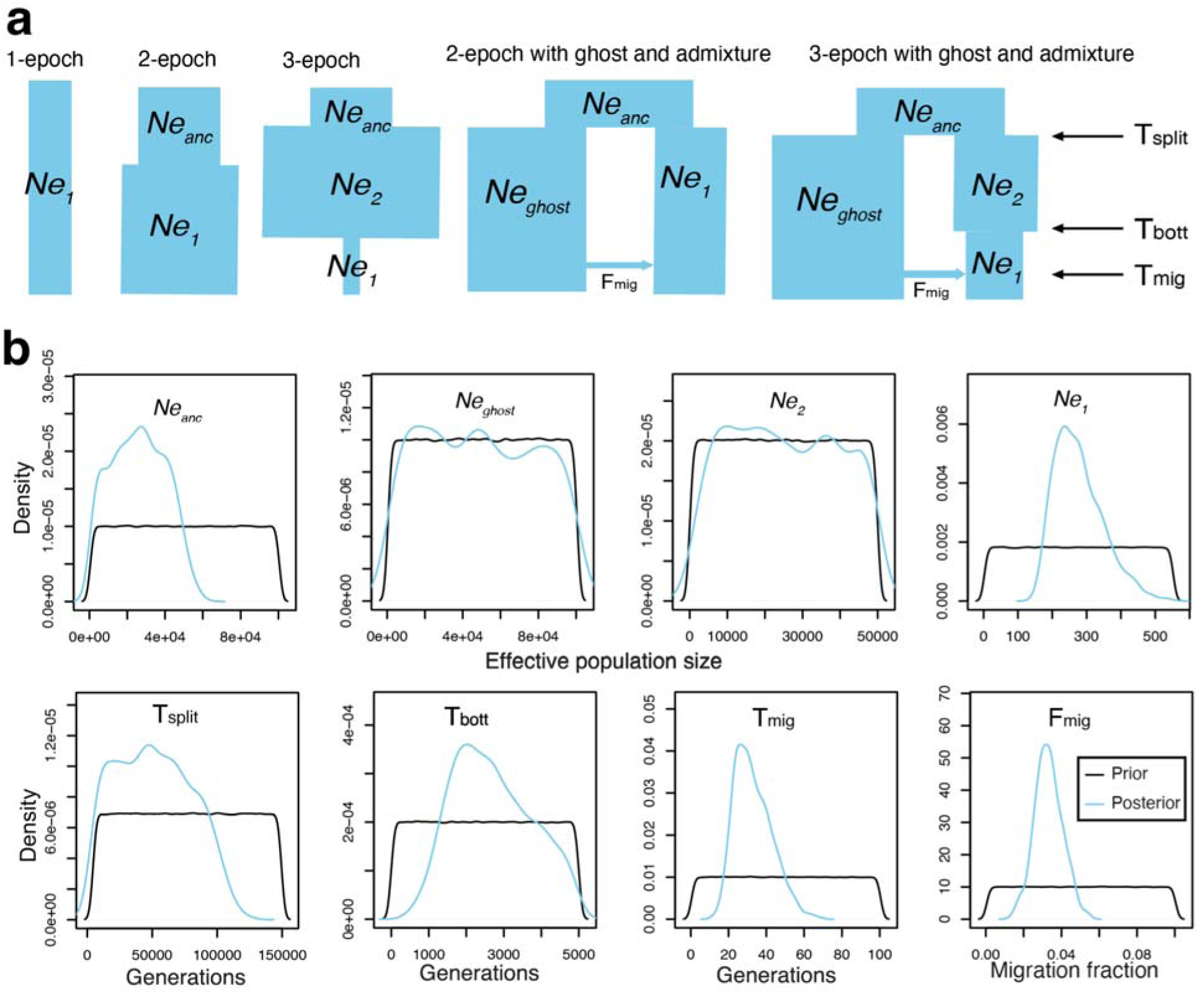
Demographic models and results for ABC inference. **a,** Pictorial representations of models tested in demographic inference (not to scale). **B,** Plots showing prior and posterior distributions of parameters for the “3-epoch ghost-admixture with a recent population decline” model from ABC. Posterior distributions are derived from the 1,000 best-fitting models out of 1 million total. Note that the *N*_e1_, T_mig_, and F_mig_ parameters are well resolved. The *N*_eanc_, T_split_, and T_bott_ parameters are moderately resolved, and the *N*_eghost_ and *N*_e2_ parameters are not resolved.

**Supplemental Fig 4.**
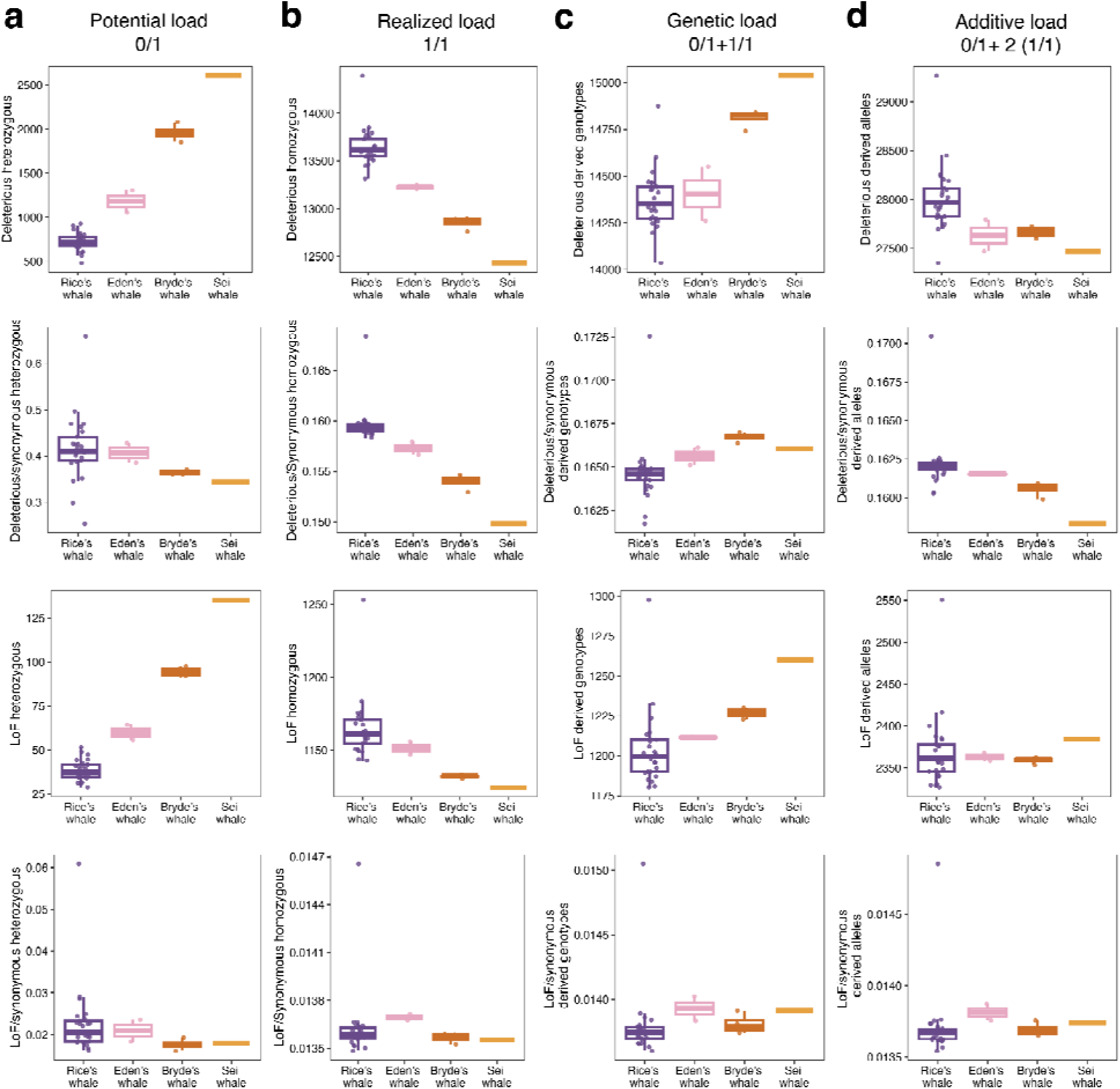
Deleterious and loss-of-function variant loads in baleen whales (extended). Deleterious (rows 1–2) and loss-of-function (LoF; rows 3–4) variant counts and ratios relative to synonymous variants are shown across four load categories in Rice’s, Eden’s, Bryde’s, and sei whales. **a,** Potential load, calculated as the total number of heterozygous (0/1) derived variants. **b,** Realized load, calculated as the number of homozygous derived (1/1) variants. **c,** Genetic load, calculated as the combined count of heterozygous and homozygous derived variants (0/1) + (1/1). **d,** Additive load, calculated as the sum of all alternative alleles across individuals. Rows 1 and 3 show raw variant counts for deleterious and LoF variants, respectively. Rows 2 and 4 show the corresponding ratios normalized to synonymous variants.

**Supplemental Fig. 5.**
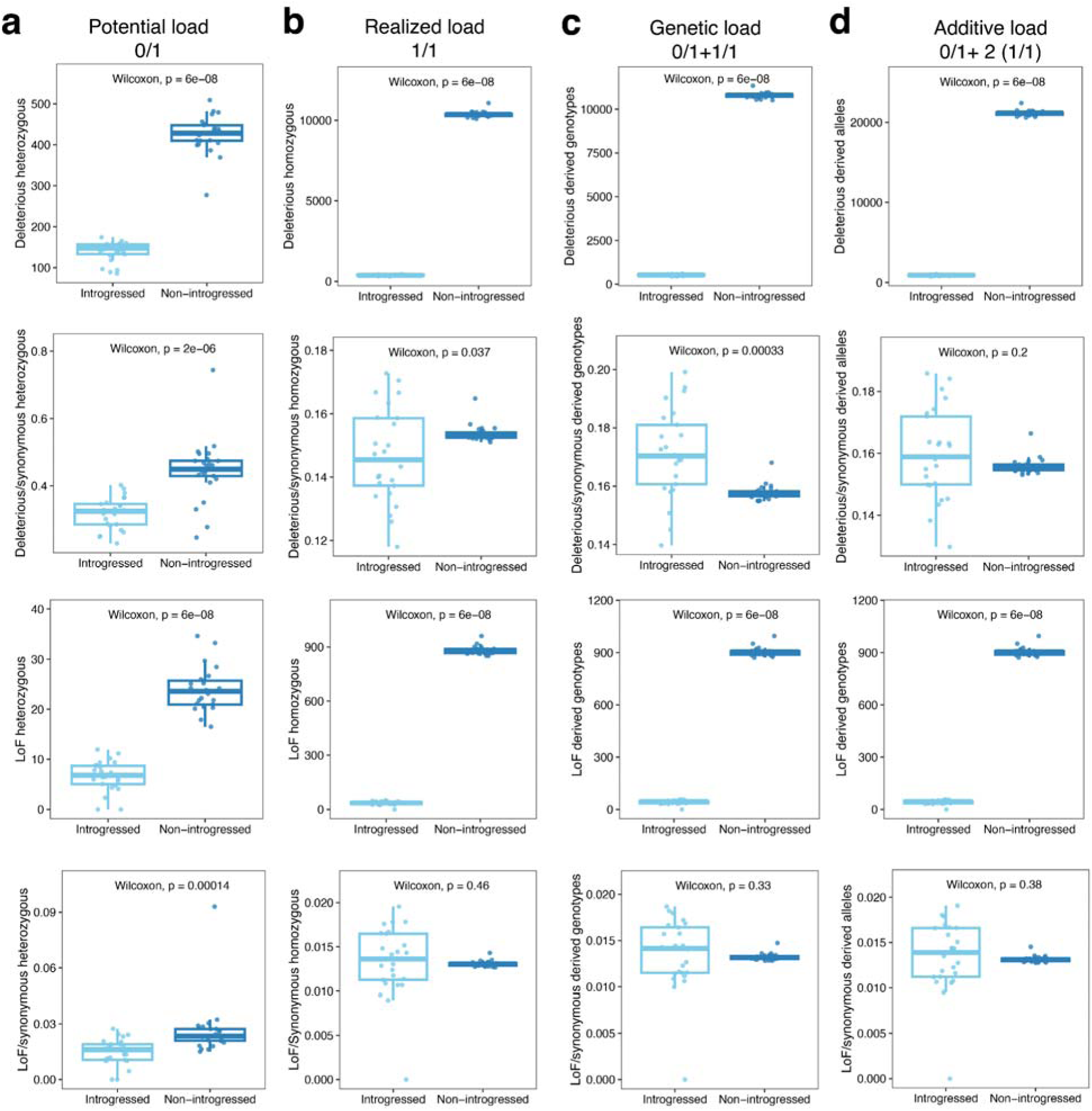
Deleterious and loss-of-function variant loads in introgressed and non-introgressed regions of the Rice’s whale genome (extended). Deleterious (rows 1–2) and loss-of-function (LoF; rows 3–4) variant counts and ratios relative to synonymous variants are shown across four load categories, partitioned by genomic context (introgressed vs. non-introgressed regions). **a,** Potential load, calculated as the total number of heterozygous (0/1) derived variants. **b,** Realized load, calculated as the number of homozygous derived (1/1) variants. **c,** Genetic load, calculated as the combined count of heterozygous and homozygous derived variants (0/1) + (1/1). **d,** Additive load, calculated as the sum of all alternative alleles across individuals. Rows 1 and 3 show raw variant counts for deleterious and LoF variants, respectively. Rows 2 and 4 show the corresponding ratios normalized to synonymous variants.

**Supplemental Fig 6.**
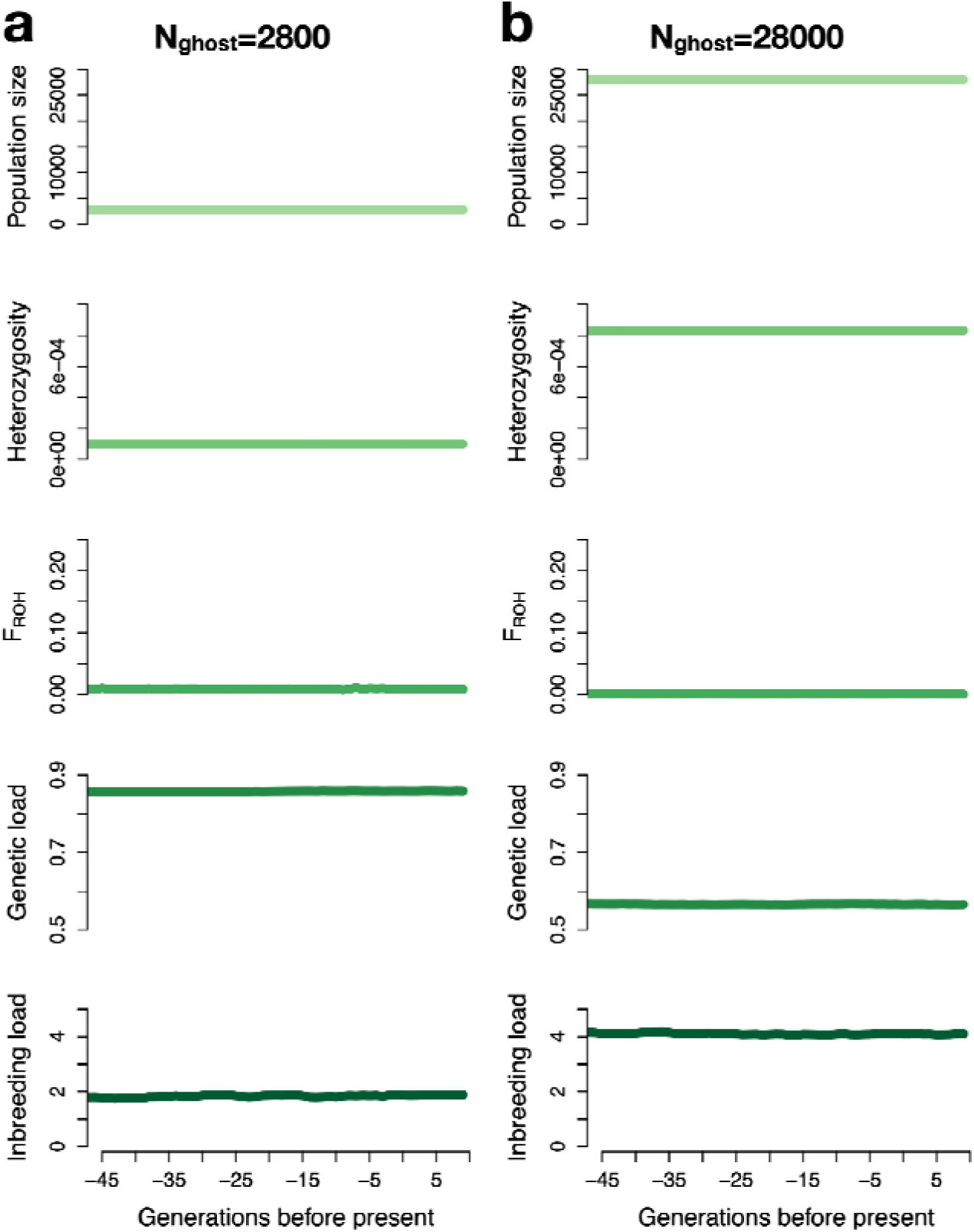
Simulation results for the ghost population when assuming N_ghost_=2800 or N_ghost_=28000. For each simulated scenario, results are shown for 45 generations before present and projected 10 generations into the future. Top panel shows effective population size over time for the ghost population, second panel shows average observed heterozygosity, third panel shows average F_ROH_ for ROH>1Mb, fourth panel shows realized genetic load, and bottom panel shows inbreeding load calculated as the diploid number of lethal equivalents.

